# SGRN: A Cas12a-driven Synthetic Gene Regulatory Network System

**DOI:** 10.1101/2023.05.08.539911

**Authors:** HyunJin Kang, John C. Fitch, Reeba P. Varghese, Curtis A. Thorne, Darren A. Cusanovich

## Abstract

Gene regulatory networks, which control gene expression patterns in development and in response to stimuli, use regulatory logic modules to coordinate inputs and outputs. One example of a regulatory logic module is the gene regulatory cascade (GRC), where a series of transcription factor genes turn on in order. Synthetic biologists have derived artificial systems that encode regulatory rules, including GRCs. Furthermore, the development of single-cell approaches has enabled the discovery of gene regulatory modules in a variety of experimental settings. However, the tools available for validating these observations remain limited. Based on a synthetic GRC using DNA cutting-defective Cas9 (dCas9), we designed and implemented an alternative synthetic GRC utilizing DNA cutting-defective Cas12a (dCas12a). Comparing the ability of these two systems to express a fluorescent reporter, the dCas9 system was initially more active, while the dCas12a system was more streamlined. Investigating the influence of individual components of the systems identified nuclear localization as a major driver of differences in activity. Improving nuclear localization for the dCas12a system resulted in 1.5-fold more reporter-positive cells and a 15-fold increase in reporter intensity relative to the dCas9 system. We call this optimized system the “Synthetic Gene Regulatory Network” (SGRN, pronounced “sojourn”).

## Introduction

Both eukaryotic and prokaryotic cells rely on their genomes to encode regulated responses to specific stimuli. In addition, multicellular organisms use the same static genome sequence in varied ways to bring about the differentiation of distinct cell types across the body. At least some of this variation is orchestrated by specific non-protein-coding sequences that interact with transcription factors (TFs) to impose robust regulatory logical circuits. One such regulatory logic is known as a gene regulatory cascade (GRC), in which suites of genes are expressed in a defined temporal ordering based on the sequential and hierarchical activation of transcriptional regulators (e.g. [1–3]).

From a synthetic biology perspective, much research has aimed at implementing synthetic regulatory systems (including GRCs) to drive novel expression patterns, both to test the accuracy of inferred models (e.g. [4]) and to drive wholly synthetic expression patterns (e.g.[5]). The advent of single-cell genomics has further underscored the need for experimental systems to test the effects of GRCs, as multiple methods have been developed to infer the sequence of transcriptional activation in GRCs at transcriptomic scale (e.g. [6,7]). Unfortunately, similarly scalable validation methods are lacking. Several modern synthetic biology efforts have implemented systems using either the clustered regularly interspaced short palindromic repeats (CRISPR)/CRISPR associated (Cas9) [5] or a combination of Cas9 and recombinases [8] to drive synthetic expression of target genes in a specified order with flexibility and scalability. Both the Cas9 and Cas9/recombinase systems used a DNase-defective Cas9 (dCas9) that had been fused to a transcriptional activation domain - VP64 (based on the herpes simplex virus protein VP16 [9]) and VPR (which further modifies VP64 to include p65 and Rta [10]), respectively. In addition, these systems also require a 3’ RNA triple helix structure from the *MALAT1* gene (triplex) to stabilize non-polyadenylated RNA. Finally, the systems use the Csy4 protein, which allows trans-activating CRISPR RNA (tracrRNA) and crisprRNA (crRNA) elements, the combination of which is referred to as a gRNA (for “guide RNA”), to be released from transcripts expressed downstream of RNA polymerase II promoters.

While these systems are quite flexible and multiplexable, we reasoned that implementing dCas12a (formerly known as Cpf1) might provide certain advantages over dCas9. The dCas9 system requires two accessory factors for full functionality: a 75-100 nucleotide (nt) long tracrRNA [11], and the Csy4 enzyme (or ribozymes [12]) for processing gRNAs from longer RNAs through the ability of Csy4 to cleave RNAs at 28 nt recognition sites. In contrast, Cas12a does not require a tracrRNA and can process its own guiding crRNA from longer transcripts when the guide is flanked by 20 nt direct repeat (DR) sequences [13], potentiating a more streamlined system. Furthermore, Cas12a has been reported to have higher target binding specificity [14].

Based on these advantages, we sought to modify the existing dCas9-based system to be driven by dCas12a instead. We evaluated how various parameters of these two systems influenced their ability to drive expression of fluorescent reporters, including the number of target sites (TS), the length of guide RNAs, and the use of different promoters and activation domains. While the dCas9 system was initially more efficient, we observed a deficiency in nuclear localization for the previously published dCas12a activator. Replacement of the nuclear localization signal (NLS) with the one used in the dCas9 system caused the dCas12a system to greatly outperform dCas9. In fact, the difference in performance became more evident as the complexity of the cascade increased (from a two-level cascade to a three-level cascade). We call this optimized Cas12a system a Synthetic Gene Regulatory Network (“SGRN”, pronounced “sojourn”, which means “a temporary stay somewhere”).

## Materials and Methods

### Construction of plasmids

Unless otherwise noted, plasmids used in this work (**Table S1**) were obtained from Addgene. All primers were synthesized by IDT with standard desalting, except for P2F_2nd and P2R_2nd which were ordered from Operon. General PCR for cloning was performed using the NEBNext PCR master mix (NEB cat no. M0541L), with 500 nM of forward and reverse primers, 10 ng of template plasmid with the following cycling conditions on a Bio-rad C1000 Touch thermal cycler: 98 °C for 30 sec, followed by 25 cycles of 98 °C for 10 sec, 58∼76 °C for 20 sec (exact temperature for each primer pair listed in **Table S1**), and 72 °C for 30 sec/kb, and final extension at 72 °C for 2 min. In some instances, overlapping primer pairs were hybridized and amplified at a concentration of 1 μM for both the forward and reverse primer to make short dsDNA inserts. PCR products were verified with agarose gel electrophoresis or polyacrylamide TBE gel electrophoresis and isolated by gel extraction kit (Thermo Fisher cat no. K0692) as necessary. Gibson assembly was done using NEBuilder® HiFi DNA Assembly Master Mix (NEB cat no. E2621S) according to manufacturer instructions. Restriction enzymes were purchased from Thermo Scientific and NEB. Quick ligase was purchased from NEB (cat no. M2200L). The suppliers and catalog numbers of the enzymes are listed in the supplementary file. NEB5-alpha (NEB cat no. C2987I) competent *E. coli* were used for cloning plasmids smaller than 10 kb and NEBStable (NEB cat no. C3040I) competent *E. coli* were used for cloning plasmids larger than 10 kb. Plasmids were purified using a Zymo mini-prep kit (RPI cat no. ZD4210) and the concentration was determined using a NanoDrop One spectrophotometer (Thermo Fisher). We verified the sequences of the plasmids with Sanger sequencing by the University of Arizona Genetics Core (UAGC) or whole-plasmid sequencing by Primordium Lab (https://primordiumlabs.com/).

To generate a DNase-inactive Cas12a (dCas12a) construct, Glu993 of Cas12a in pCE059-SiT-Cas12a-[Activ]-M13(Addgene cat no. 128136) was mutated into alanine [27] by two rounds of overlap extension PCR [28]. A region of the Cas12a construct containing AccI and Eco91I recognition sites was then amplified in two fragments with a mutated overlap. Isolated fragments were combined into one piece by a second PCR and then assembled into AccI- and Eco91I– restricted pCE059-SiT-dCas12a-[Activ]-M13 by Gibson assembly (All primers listed in **Table S1**). Assembled products were transformed into NEBstable competent cells.

#### Constructs relevant to Figure 1

To create dCas12a-crRNA constructs with the P1 crRNA, P2 crRNA, and P3 crRNA, DNA fragments containing the 20 bp DR sequences and crRNA1, crRNA2, or crRNA3 for dCas12a were amplified by primer dimer extension PCR and inserted into the SapI-restricted pCE059-SiT-dCas12a-[Activ]-M13 by Gibson assembly. The gRNA constructs with 28 bp Csy4 recognition sites and gRNAs for dCas9 were built as follows: the EF1a-triplex-28-gRNA1-28 plasmid was built through 2 steps. First, inverse PCR from CMVp-dsRed2-Triplex-28-gRNA1-28 (Addgene cat no. 55200) using a pair of primers including SacI restriction sites generated a linear plasmid, excluding dsRed2, followed by SacI restriction and ligation. Second, the CMV promoter of the dsRed2-free plasmid was replaced with the EF1α promoter amplified from pCEP4_WT_OCT4 (Addgene cat no. 40629) by Gibson assembly after KpnI and SacI restriction. For the EF1a-triplex-28-gRNA2-28 and EF1a-triplex-28-gRNA3-28 plasmids, amplicons of gRNA2 and gRNA3 were amplified using primers that created XbaI and NheI sites at the ends and then gRNA1 of the EF1a-triplex-28-gRNA1-28 plasmid was replaced by traditional restriction-ligation cloning after digesting the plasmid with XbaI and NheI as well (all primers listed in **Table S1**).

**Figure 1.**
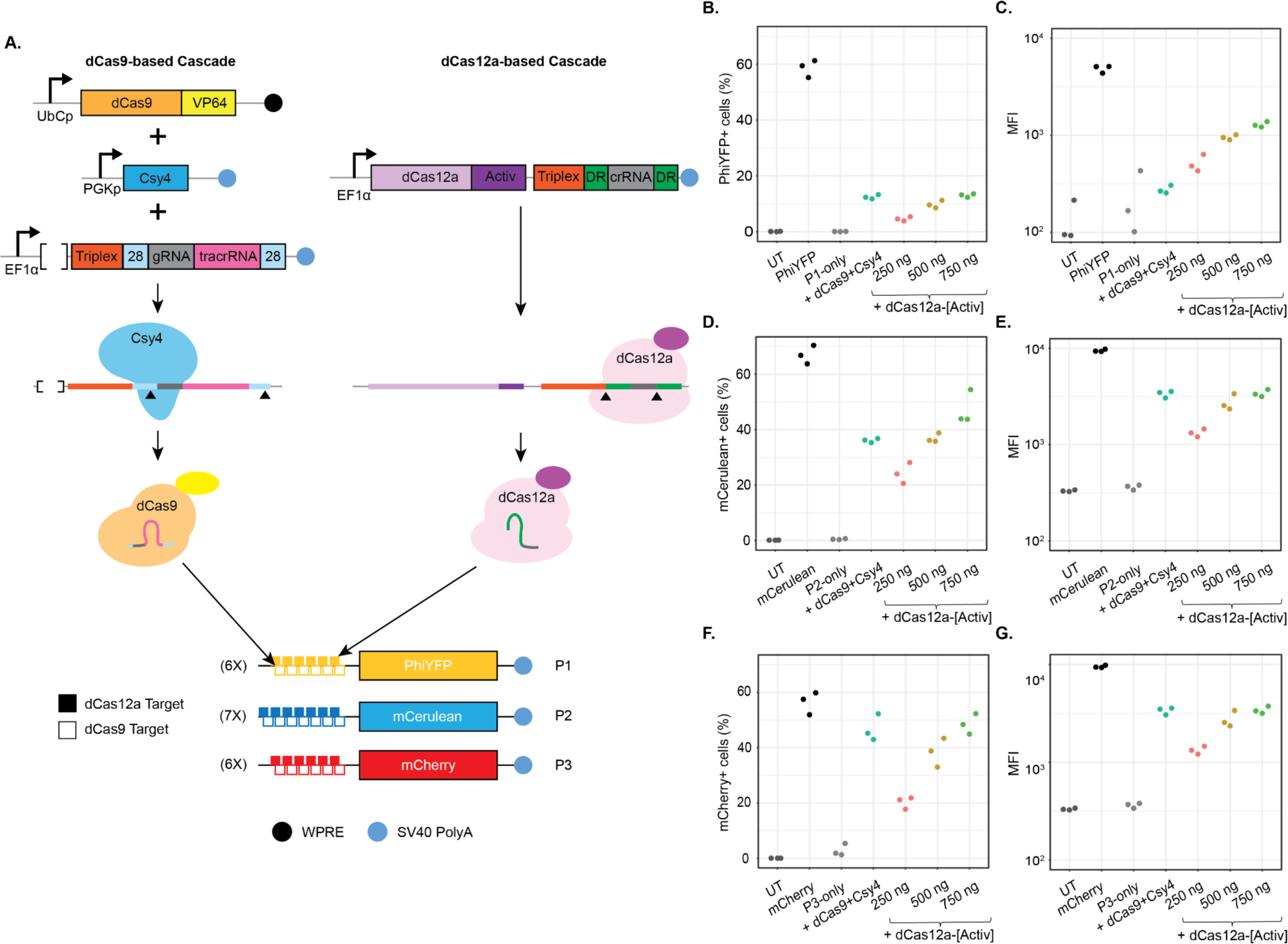
Initial implementation of SGRN, a dCas12a-based transcriptional cascade. (**A**) Schematic of two transcriptional cascades. The left-hand side depicts the components of a previously developed dCas9-based transcriptional cascade [5]. This includes a dCas9-VP64 fusion, the Csy4 gene, and the first transcript in the cascade - a non-coding transcript that includes gRNA1, which guides dCas9-VP64 to P1. Csy4 cleaves the gRNA-containing transcript at 28 nt recognition motifs and the triplex stabilizes the resulting non-polyadenylated transcript. The right-hand side depicts the components of our dCas12a-based transcriptional cascade. The three components required in the dCas9 system are replaced by a single unit that encodes dCas12a, which can process its own crRNAs and therefore includes a crRNA targeting P1 in its own 3’ UTR. dCas12a cleaves the crRNA at the DRs and the triplex stabilizes the resulting non-polyadenylated transcript. (**B-G**) Quantification of the activity of the cascade system on three different promoters, measuring both %positive for the fluorescent reporter (**B,D,F**) and the MFI (on Log_10_ scale) for positive cells (**C,E,G**). (**B,C**) Activity of the cascades in targeting the P1 construct which includes the PhiYFP reporter gene. (**D,E**) Activity of the cascades when targeting P2 using the mCerulean reporter gene. (**F,G**) Activity of the cascades targeting P3 and using the mCherry reporter gene. In all cases, 250ng of dCas9-VP64 transfected was compared to transfection with 250, 500, or 750 ng of the dCas12a-[Activ] construct. “UT” denotes “untransfected”, “PhiYFP”, “mCerulean”, and “mCherry” denote constitutively expressed reporters that use the CMV promoter. To assess background transcription from the reporter genes, the downstream target plasmid was transfected without the upstream activator for each reporter - denoted as “P1-only”, “P2-only”, or “P3-only”, respectively.

The P3-mCherry-pA plasmid was initially constructed to contain 7 target sites for dCas9 (without consideration of dCas12a sites). To build this construct, we performed Golden Gate cloning using BsmBI, a type IIS restriction enzyme. P2-mCerulean-pA was digested with NcoI and NheI to remove the P2 promoter region. Forward (P3_BsmBI_fwd) and reverse (P3_BsmBI_rev) oligomers (**Table S1**) were annealed in a buffer containing 10mM Tris-HCl (pH7.5) and 50 mM NaCl to create a 1xTS construct with a BsmBI-v2 recognition site and cohesive ends compatible with the NheI and NcoI-restricted P2 plasmid. Ligated products were transformed with the NEB5-alpha strain. After verifying the sequence of the 1xTS and the BsmBI site of the new plasmid by Sanger sequencing, the plasmid was restricted with BsmBI-v2 to repeat inserting the annealed oligo to add another TS and and a new BsmBI-v2. After repeating this cloning for a total of 7xTS, we ligated annealed synthetic dsDNA carrying the original P2 5’-UTR into the BsmBI-v2-treated P3-7xTS-BsmBI-pA to remove the temporary BsmBI-v2 site and to fix the 5’-UTR. We then replaced the mCerulean gene carried in the new P3 plasmid first with the dsRed2 gene amplified from the CMVp-dsRed2-Triplex-28-gRNA1-28 (Addgene cat no. 55200) into the FseI-treated P3-7xTS by restriction-ligation to create P3-7xTS-dsRed2. After determining that dsRed2 was not ideal for a multi-FP panel, the P3-7xTS-dsRed2 plasmid was digested with AgeI and SacI to replace the dsRed2 with mCherry amplified from mCherry-c2 (kind gift of Dr. Ghassan Mouneimne). However, this P3 construct had different numbers of target sites for dCas9 and dCas12a (7xTS for dCas9 and 6xTS for dCas12a). Therefore, we used PCR to attempt to add a 7th TS for dCas12a using a primer in the 5’UTR and a primer that could anneal to any of the dCas9 target sites and ensured a dCas12a TS on the opposite strand. After PCR amplification, the large fragments were gel-extracted and cloned them into the NheI-restricted P3-mCherry construct, hoping to isolate a product with 7xTS for both dCas9 and dCas12a. Ultimately, we identified a colony that only had 6xTS for both dCas9 and dCas12a (verified by Sanger sequencing). Since the number of targets was now balanced between the two enzymes, we decided to proceed with this construct.

#### Constructs relevant to Figure 2

To build the dCas12a-crRNA2(16 nt) and dCas12a-crRNA2(18 nt), the amplified crRNA2 fragments with 16 and 18 bp length and SapI-digested pCE059-SiT-dCas12a-[Activ]-M13 were subjected to Gibson Assembly. NEBstable *E. coli* was transformed with these new dCas12a plasmids. The EF1a-triplex-28-gRNA2(16 nt)-28 and EF1a-triplex-28-gRNA2(18 nt)-28 constructs for dCas9 were built by overlap extension PCR of DNA fragments carrying 16 or 18 bp gRNA sequences to include XbaI and NheI restriction sites at the ends. After restriction digest and gel extraction of the plasmid backbone and inserts, the new guides were ligated into the construct and the product was transformed into NEB5-alpha competent *E. coli*.

**Figure 2.**
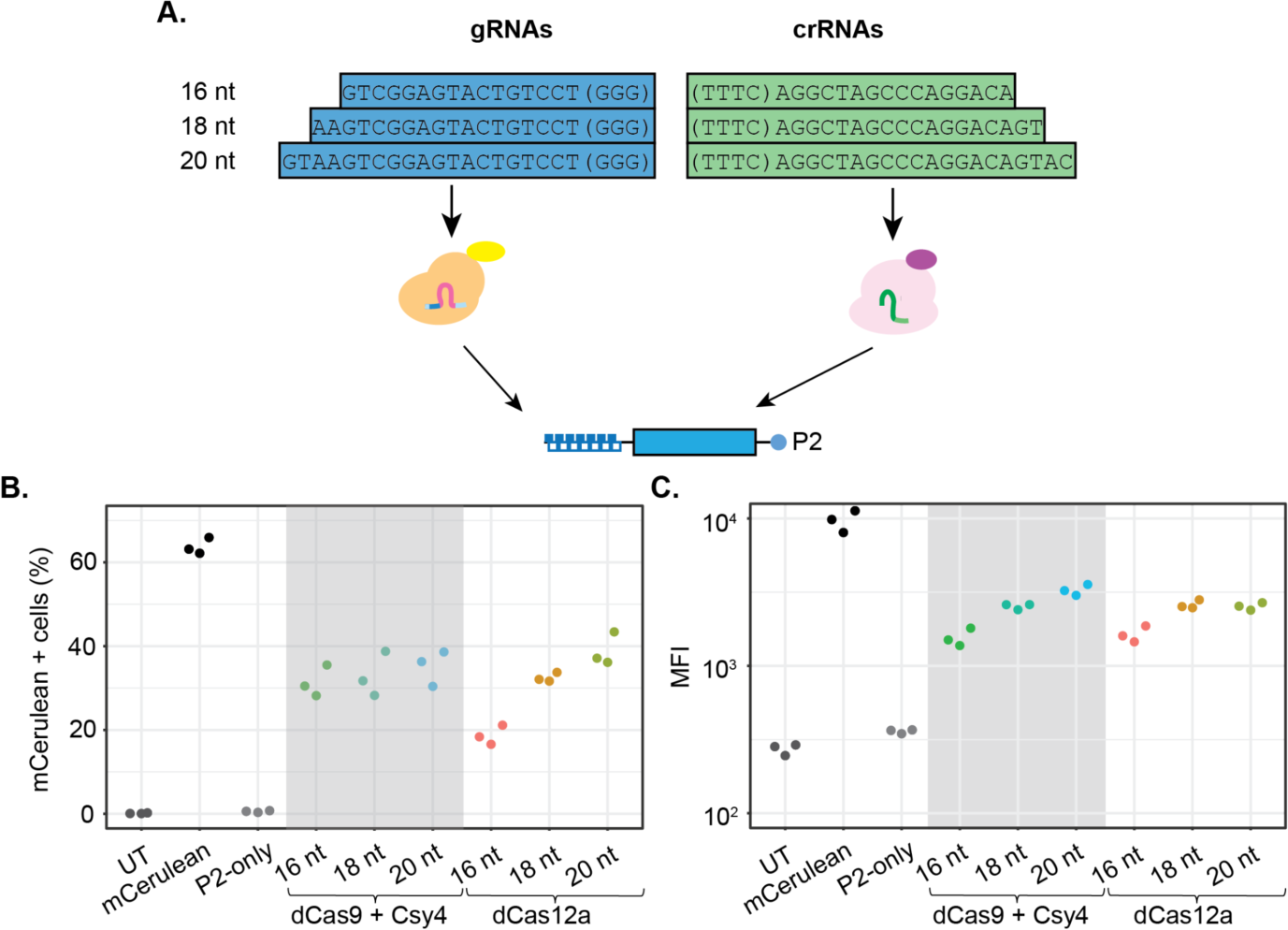
Effect of guide RNA length on reporter activity. (**A**) Schematic of the experiment testing different lengths of guide RNAs for both dCas9 and dCas12a. Guides of length 16-20 nt (in 2nt increments) were designed. (**B**) %positive measurement of P2 targeted with different length guides. Y-axis indicates the percent of cells detected as positive for mCerulean (downstream of P2). (**C**) MFI (on Log_10_ scale) for positive cells in each condition. “UT” denotes untransfected control cells. “mCerulean” is a constitutively active control plasmid. “P2-only” includes the target reporter, but not the upstream Cas activator or targeting guide RNA. The remaining samples indicate the guide length used (16-20 nt) and the activating Cas enzyme (dCas9 with Csy4 for processing gRNAs or dCas12a alone).

#### Constructs relevant to Figure 3

The P2-mCerulean-pA plasmids with 1 to 7 target sites in the promoter region were generated by Golden Gate Assembly using BsmBI-v2 in the same way described for the P3-mCherry-pA construct above. Briefly, the original promoter region of the P2-mCerulean-pA plasmid was removed by digestion with NheI and NcoI and inserted a dsDNA fragment with a BsmBI site (BsmBI-mCerulean-pA), followed by BsmBI restriction and insertion of another dsDNA fragment containing a 1xTS and a new BsmBI site to generate P2-1xTS-BsmBI-mCerulean. After Sanger sequencing, the verified construct was restricted with BsmBI-v2 and ligated with an additional fragment containing an additional target site and a new BsmBI restriction site. After creating all constructs with the desired number of target sites (up to 7), the 5’ UTR was corrected for each by inserting annealed oligonucleotides with correct sequences and all constructs were verified by Sanger sequencing.

**Figure 3.**
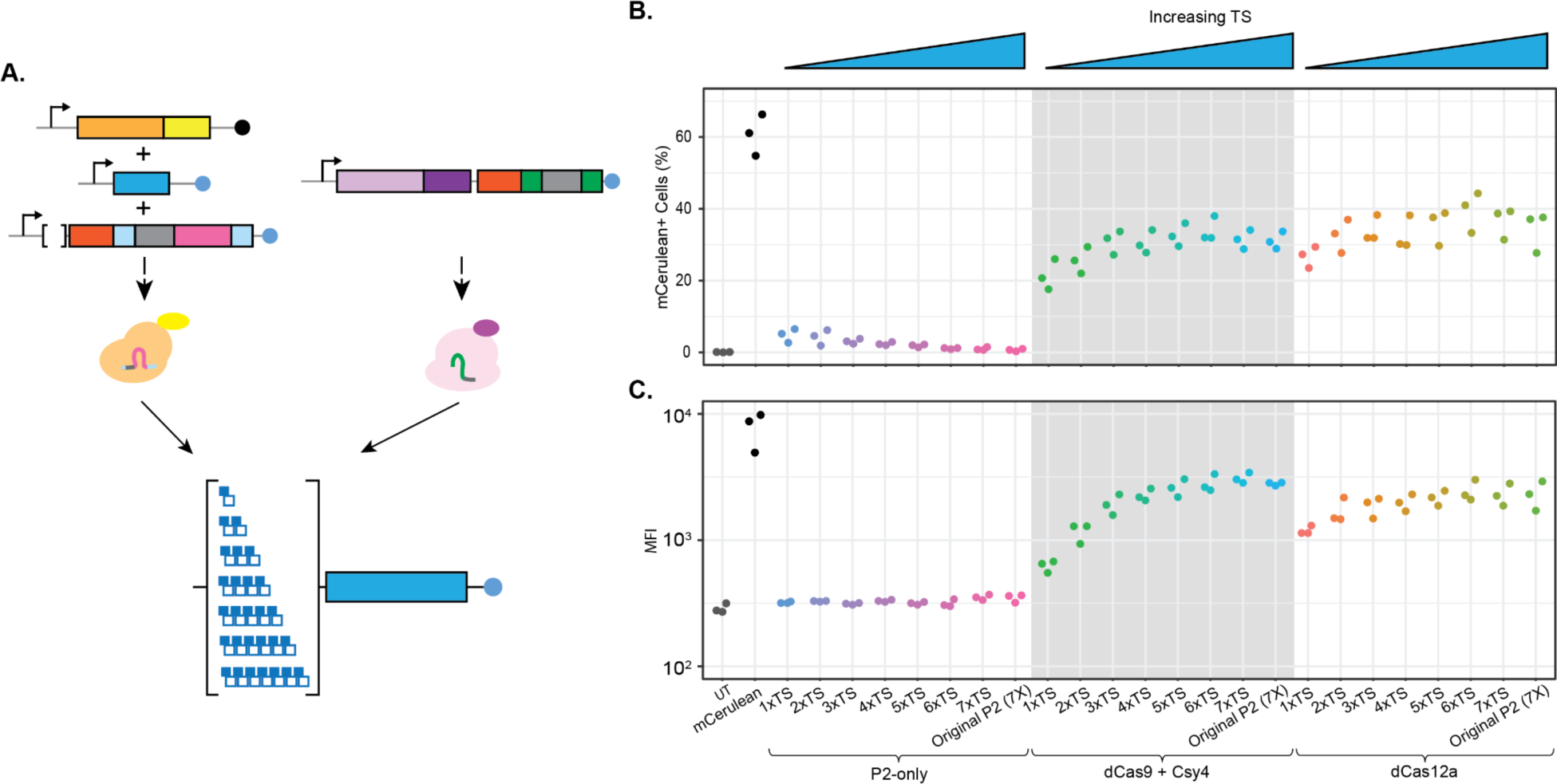
Effect of number of target sites in synthetic promoter on reporter activation. (**A**) Schematic of experiment design. Either dCas9 and Csy4 or dCas12a were used to target a series of synthetic promoters with differing numbers of TSs (from one to seven). (**B**) %positive measurement of P2 targeted with different numbers of TSs in the promoter. Y-axis indicates the %positive for mCerulean (downstream of P2). (**C**) MFI (on Log_10_ scale) for positive cells in each condition. “UT” denotes untransfected control cells. “mCerulean” is a constitutively active control plasmid. The first series of target plasmids bracketed by “P2-only” includes the target reporter, but not the upstream Cas activator or targeting guide RNA. The subsequent two series of promoters were targeted by either dCas9 and Csy4 or dCas12a (noted by the brackets). The “Original P2” construct was used in the previous experiments and is included here for consistency due to slight differences in the promoter structure.

#### Constructs relevant to Figure 4

To build the UbCp-dCas12a-[Activ]-crRNA2, the amplified UbC promoter region from the dCas9-VP64 replaced the EF1α promoter of the EF1a-dCas12a-[Activ]-crRNA2 by traditional restriction-ligation using PacI and KpnI restriction enzymes. The EF1a-dCas12a-VP64-crRNA2 was generated by Gibson assembly, replacing the [Activ] domain with VP64 amplified from the dCas9-VP64. The EF1a-dCas12a-VP64-crRNA1 and EF1a-dCas12a-VP64-crRNA3 constructs were generated by replacing the MALAT1 3’ triplex–DR-crRNA2 region in the EF1a-dCas12a-VP64-crRNA2 with amplified MALAT1 3’ triplex-DR-crRNA1 or MALAT1 3’ triplex-DR-crRNA3 from the dCas12a-[Activ]-crRNA1 and dCas12a-[Activ]-crRNA3, respectively, by Gibson assembly.

**Figure 4.**
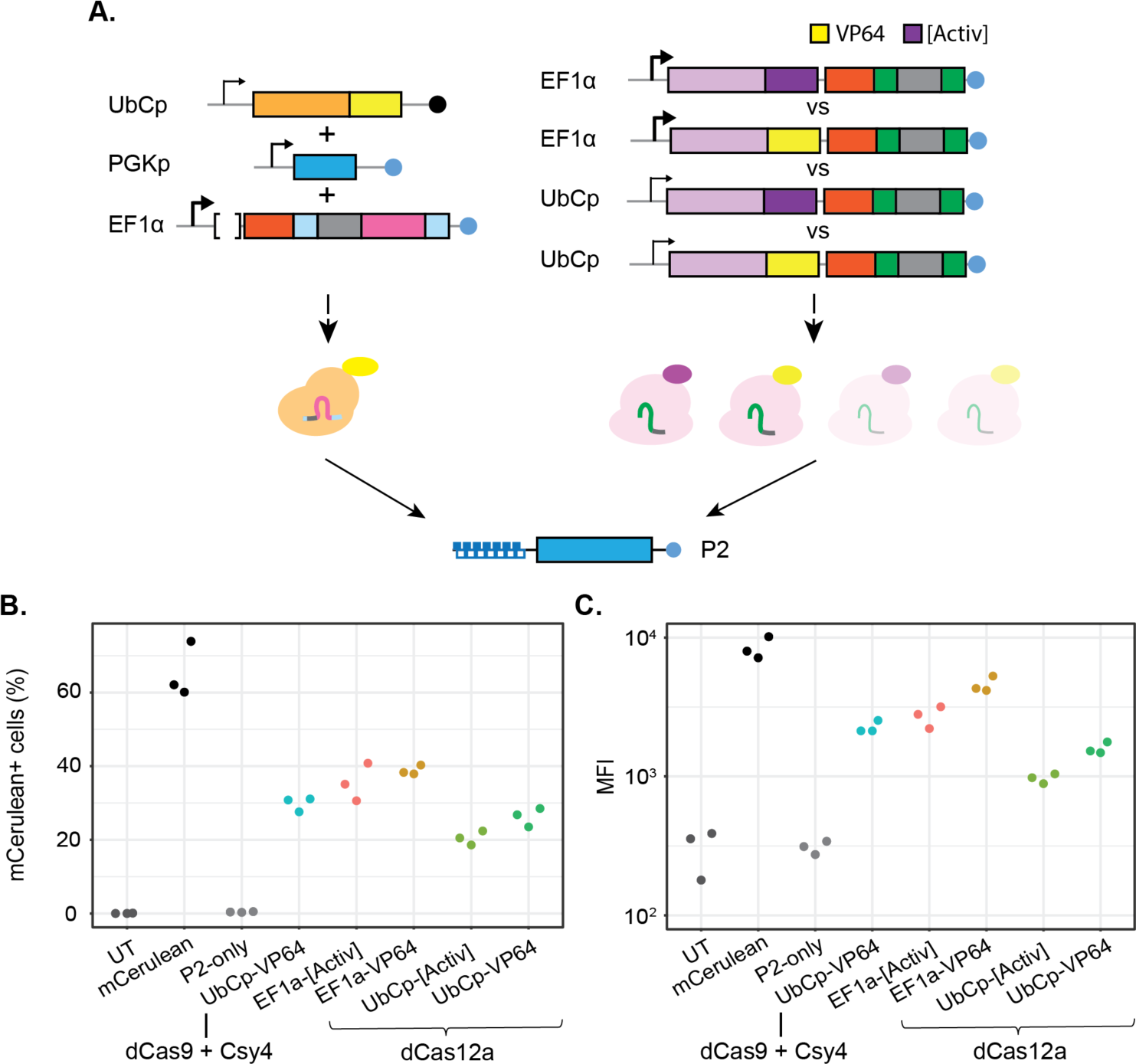
Effect of promoter and activation domain on cascade activity. (**A**) Schematic of experimental design. The original dCas9 and Csy4 system was compared to dCas12a systems that used different constitutive promoters (EF1α versus UbCp) or different transcriptional activation domains (VP64 versus [Activ]). (**B**) Percent positive measurement of P2 targeted with different activators. Y-axis indicates the percent of cells detected as positive for mCerulean (downstream of P2). (**C**) MFI values (on Log_10_ scale) for positive cells in each condition. “UT” denotes untransfected control cells. “mCerulean” is a constitutively active control plasmid. “P2-only” includes the target reporter, but not the upstream Cas activator or targeting guide RNA. The remaining conditions indicate the Cas enzyme, the promoter used for expression of the Cas enzyme and the transcriptional activation domain fused to the Cas enzyme.

#### Constructs relevant to Figure 5

To generate GFP-fused dCas reporter constructs, we started with a version of dCas9-3xNLS-VP64 that had been moved to the pGH125 backbone - replacing the original dCas9-NLS of pGH125_dCas9-Blast (Addgene cat no. 85417) with dCas9-3XNLS-VP64 (Addgene cat no. 55195) by Gibson assembly. dCas9 (pGH125-GFP-dCas9-3xNLS-VP64) was generated by inserting GFP amplified from EX-EGFP (kind gift of Dr. Casey Romanoski) with primers incorporating BsiWI sites into BsiWI-restricted pGH125-dCas9-3xNLS-VP64. In the same way, pGH125-GFP-dCas12a-1xNLS-Activ, pGH125-GFP-dCas12a-3xNLS-Activ, pGH125-GFP-dCas12a-3xNLS-VP64 were created by combining amplified DNA fragments of GFP, dCas12a, relevant NLS and TAD with restricted pGH125-GFP-dCas9-3xNLS-VP64 as a backbone by Gibson assembly (further described in **Table S1**).

**Figure 5.**
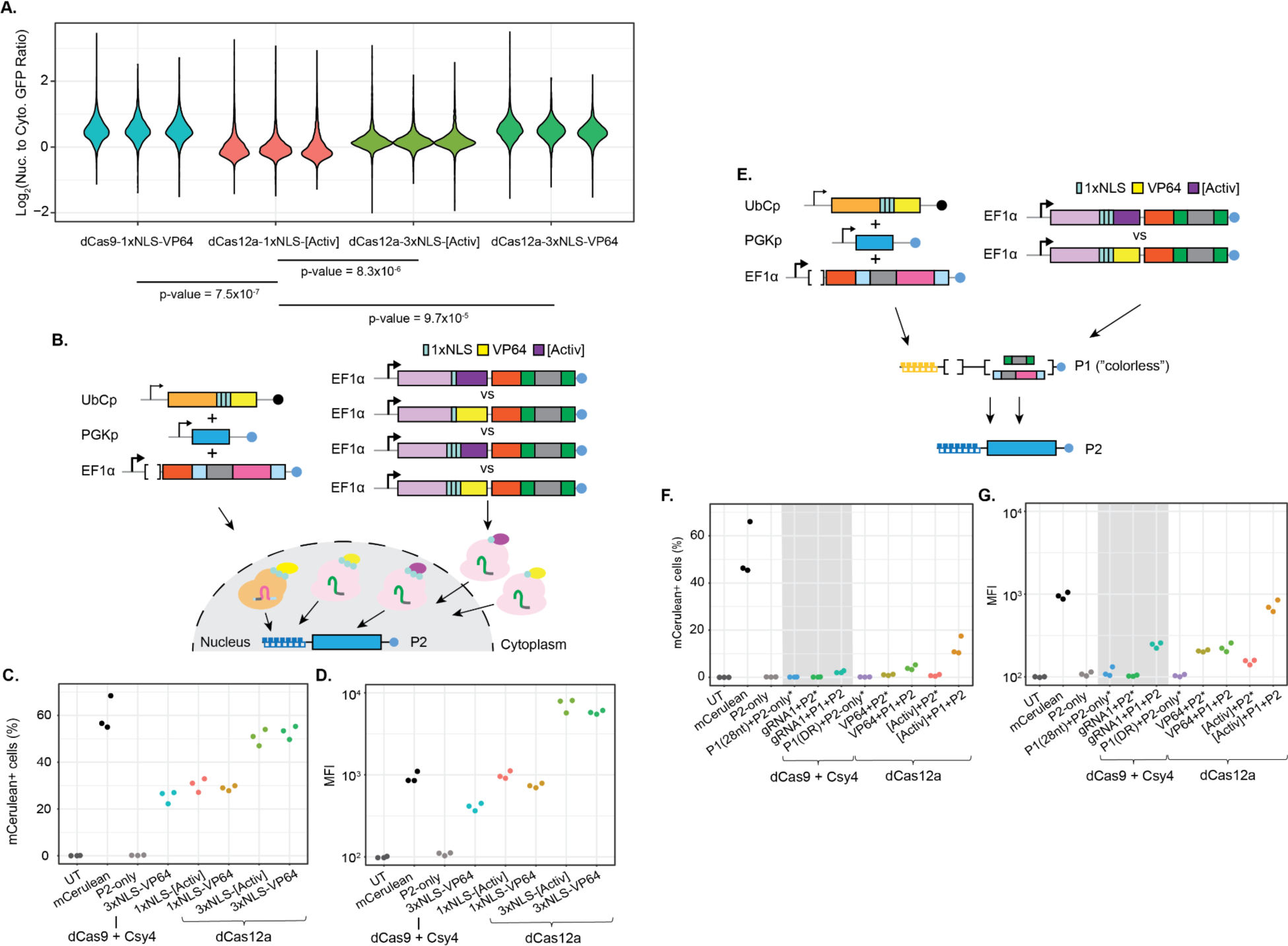
Effect of nuclear localization on SGRN activity. (**A**) Violin plots of Log_2_(Nuclear/Cytoplasmic) GFP intensity ratios from fluorescence microscopy. Each violin represents ratios for all individual cells measured. All transfections were performed in triplicate. The p-values below the x-axis indicate that all three alternative constructs are more nuclear than the original dCas12a-[Activ]. (**B**) Schematic of different SGRN NLS variants tested. The original dCas9 system (with a 3xNLS) was tested against the initial SGRN system (1xNLS), as well as variants of SGRN that contained 3xNLS domains. (**C**) %positive cells for mCerulean when targeting the P2 reporter construct with the different activators. (**D**) MFI values (on Log_10_ scale) for the different constructs tested. (**E**) Schematic of our three–level cascade design, in which the initial guide RNA is expressed from a constitutive promoter (EF1α). It guides the Cas systems to activate the second level, P1 (“colorless”), which also expresses the third guide RNA to activate P2. (**F**) %positive cells for mCerulean (the final stage in the cascade) when targeting the cascade with the different activators. (**G**) MFI values (on Log_10_ scale) for the different constructs tested. “UT” denotes untransfected control cells. “mCerulean” is a constitutively active control plasmid. “P2-only” includes the target reporter, but not the upstream Cas activator or targeting guide RNA. (**C-D**) The remaining conditions indicate the Cas enzyme, the number of NLS sequences fused to the Cas enzyme, and the transcriptional activation domain fused to the Cas enzyme. (**F-G**) The remaining conditions include negative controls (“P1(28nt)+P2-only”, “gRNA1+P2-only”, “P1(DR)+P2-only”, “VP64+P2-only”, “[Activ]+P2-only” - all marked by an “*”) and the fully functional three-level cascades with the different activators (“gRNA1+P1+P2”, “VP64+P1+P2”, “[Activ]+P1+P2”).

dCas12a-3xNLS-[Activ]-crRNA2 was generated by Gibson assembly - combining amplified fragments containing a portion of dCas12a, a 3xNLS, and an [Activ] domain with Eco91I- and BamHI-restricted dCas12a-[Activ]-crRNA2. This new plasmid was also used as a backbone to build dCas12a-3xNLS-VP64-crRNA2 by replacing the 3xNLS-[Activ] with amplified 3xNLS-VP64 from pGH125-GFP-dCas9-3xNLS-VP64 by Gibson assembly.

In order to generate dCas12a-3xNLS-[Activ]-crRNA1-DR and dCas12a-3xNLS-VP64-crRNA1 for the three-level cascade, a crRNA1-containing portion of dCas12a-[Activ]-crRNA1 was amplified and ligated into NdeI- and XbaI-restricted dCas12a-3xNLS-[Activ]-crRNA2 and dCas12a-3xNLS-VP64-crRNA2, respectively. The colorless P1-triplex-DR-crRNA2-DR for dCas12a and P1-triplex-28-gRNA2-28 for dCas9 were created by inverse PCR to exclude PhiYFP from P1-PhiYFP-DR-crRNA2-DR and P1-PhiYFP-28-gRNA2-28 using outward facing primers sitting at the 3’-end of the 5’-UTR and the 5’-end of the 3’-UTR and including SacI restriction sites (**Table S1**). The two plasmids used as PCR templates were built in-house previously; The P1-PhiYFP-28-gRNA2-28 was generated by inserting gRNA2 amplified from CMVp-dsRed2-28-gRNA2-28 into FseI- and MfeI-restricted P1-PhiYFP-pA and P1-PhiYFP-DR-crRNA2-DR-pA were constructed by Gibson assembly, replacing the MALAT1 triplex-28-gRNA2-28 of P1-PhiYFP-28-gRNA2-28 with MALAT1 triplex-DR-crRNA2-DR amplified from dCas12a-[Activ]-crRNA2. The PCR products were subsequently restricted and ligated to circularize the construct before NEB5-alpha transformation.

#### A generic set of SGRN plasmids

To facilitate future research implementing this dCas12a system, we designed a generic set of plasmids that can be easily modified to target (and be controlled by) arbitrary guide sequences for activation. First, we generated a set of plasmids containing dCas12a-based transcriptional activators with a cloning site for introducing crRNAs in the 3’ UTR of the dCas12a gene. These include the following constructs:

- “pCE059-SiT-ddCas12a(E993A)-1xNLS-[Activ]-triplex-DR-M13-DR-pA” (Addgene ID 202042) - a DNase-dead Cas12a fused to a 1xNLS and the [Activ] transactivation domain, plus an M13 site flanked by SapI restriction sites and Cas12a DRs. The plasmid can be SapI-digested to remove the M13 sequence and hybridized oligos containing the desired crRNA can be ligated in its place.
- “pCE059-SiT-ddCas12a(E993A)-3XNLS-[Activ]-triplex-DR-M13-DR-pA” (Addgene ID 202043) - a DNase-dead Cas12a fused to a 3xNLS and the [Activ] transactivation domain, plus an M13 site flanked by SapI restriction sites and Cas12a DRs. The plasmid can be SapI-digested to remove the M13 sequence and hybridized oligos containing the desired crRNA can be ligated in its place.
- “pCE059-SiT-ddCas12a(E993A)-3XNLS-VP64-triplex-DR-M13-DR-pA” (Addgene ID 202044) - a DNase-dead Cas12a fused to a 3xNLS and the VP64 transactivation domain, plus an M13 site flanked by SapI restriction sites and Cas12a DRs. The plasmid can be SapI-digested to remove the M13 sequence and hybridized oligos containing the desired crRNA can be ligated in its place.

Next, we designed a set of plasmids that can be modified to serve as an arbitrary number of tiers of the cascade. This involved generating a set of plasmids that allow for introduction of tandem arrays of target sites in the promoter to make them Cas-responsive and (for a subset) the restriction site-flanked guide sites in the 3’ UTR were also included so that the elements can in turn activate a downstream target. The constructs were as follows:

- “Px-PhiYFP-pA” (Addgene ID 202045) - contains a cloning site in place of the promoter so that targeting arrays can be introduced upstream of the PhiYFP reporter gene. The promoter is built with a combination of SacI and BamHI digestion, iterative introduction of BsmBI site-containing target sites, and removal of the BsmBI site once enough repeats have been introduced.
- “Px-mCerulean-pA” (Addgene ID 202048) - contains a cloning site in place of the promoter so that targeting arrays can be introduced upstream of the mCerulean reporter gene. The promoter is built with a combination of SacI and BamHI digestion, iterative introduction of BsmBI site-containing target sites, and removal of the BsmBI site once enough repeats have been introduced.
- “Px-mCherry-pA” (Addgene ID 202049) - contains a cloning site in place of the promoter so that targeting arrays can be introduced upstream of the mCherry reporter gene. The promoter is built with a combination of SacI and BamHI digestion, iterative introduction of BsmBI site-containing target sites, and removal of the BsmBI site once enough repeats have been introduced.
- “Px-PhiYFP-triplex-DR-M13-DR-pA” (Addgene ID 202046) - contains a cloning site in place of the promoter so that targeting arrays can be introduced upstream of the non-coding transcript harboring a DR-flanked crRNA cloning site in the 3’ UTR. For this construct, the crRNA in the 3’ UTR is introduced first by restricting with BsmBI and ligating in annealed oligos. After introducing the crRNA, the promoter can be built with a combination of SacI and BamHI digestion, iterative introduction of BsmBI site-containing target sites, and removal of the BsmBI site once enough repeats have been introduced.
- “Px-PhiYFP-triplex-28-M13-28-pA” (Addgene ID 202047) - contains a cloning site in place of the promoter so that targeting arrays can be introduced upstream of the non-coding transcript harboring a 28 nt-flanked gRNA cloning site in the 3’ UTR. For this construct, the gRNA in the 3’ UTR is introduced first by restricting with BsmBI and ligating in annealed oligos. After introducing the gRNA, the promoter can be built with a combination of SacI and BamHI digestion, iterative introduction of BsmBI site-containing target sites, and removal of the BsmBI site once enough repeats have been introduced.

Finally, we designed a construct for the first tier of the dCas9-based cascade that is expressed from the EF1α promoter and contains a SapI-flanked cloning site in the 3’UTR for adding a gRNA to target a downstream target: “EF1a-Triplex-28-M13-28-pA” (Addgene ID 202041). Detailed protocols for modifying all of these constructs are provided on the Addgene website.

### Transient Transfections

HEK293T cells cultured in DMEM (Gibco cat no. 11995) supplemented with 10% FBS (Gibco cat no. 10437-028) and 100 U/ml Penicillin Streptomycin (Gibco cat no. 15140–122). For transfections, 1×10^5^ cells were seeded in each well of a 12-well plate and incubated overnight before transfection. The cells were transfected with plasmids using Lipofectamine 3000 (Invitrogen cat no. 3000008) according to the manufacturer’s instructions with a slight modification. The total amount of plasmid transfected per sample was 1 μg, except for the three-level cascade experiment, which used 1.25 μg of total DNA. The total amount of DNA was kept the same across all samples by adding pUC19 as necessary. For controls, we transfected 500 ng of pCDNA-PhiYFP, pCDNA-mCerulean, and/or mCherry-c2. For each sample, 250 ng of each plasmid needed was combined with pUC19 (if necessary) and 1.6 μl/μg DNA of P3000 reagent in 50 μl of Opti-MEM (Gibco cat no. 51985034). Lipofectamine 3000 (1.2 μl/μg DNA) was diluted in a separate tube before adding it to the DNA-P3000 mixture. DNA-P3000-Lipofectamine mixture was incubated at room temperature for 15 min. During the incubation, cell culture media was replaced with 500 μl of DMEM with 5% FBS and no antibiotics. After 6 h of incubation of cells with the transfection mixture, 1 ml of complete media (DMEM, 10% FBS, 100 U/ml Penicillin Streptomycin) was added, and cells were incubated for 48 h before flow cytometry analysis. The cells for imaging were transfected with 1 ug of GFP-fused dCas9 or dCas12a in the transfection reagent described above and incubated for 24 h before imaging.

### Flow cytometry

Transfected cells were dissociated with 0.25% Trypsin/EDTA (Gibco cat no. 25200056), pelleted by spinning at 500 x g for 5 min at 4°C, resuspended with 0.1% FBS in cold PBS (Gibco cat no. 10010023), and kept on ice until scanning on a BD LSRII Flow Cytometer. Unless otherwise indicated, Am-Cyan (525/50 nm peak/bandpass filter), FITC (530/30 nm), PE-Cy5 (660/20 nm) channels were used to detect mCerulean (475 nm fluorescence peak maximum), PhiYFP (537 nm peak maximum), and mCherry (610 nm peak maximum), respectively. Ten thousand single cells per sample were recorded based on gates set using the negative controls, and the percentage of fluorescence-positive cells and median fluorescence intensity (MFI) were initially measured as experimental outputs for analysis by using BD FACSDiva software (v9). Subsequently, data was re-analyzed and gates were re-set using tools available in the R programming language (see Data Analysis section below). Three biological replicates were obtained from independent transfections on different days for each experiment. In the experiments testing the effect of improved nuclear localization of dCas12a with a 3xNLS, mCerulean expression was so high that we had to use the Pacific Orange channel (560/40 nm; ∼20% peak mCerulean emission) to properly separate positive and negative cells.

### Imaging

Transfected cells were dissociated with 0.05% Trypsin/EDTA (Gibco cat no. 25300) and pelleted by spinning at 500 x g for 5 min at 4°C. Cells were resuspended in PBS and then fixed with 1% formaldehyde for 10 minutes at room temperature. After quenching the fixation with 125 mM glycine, the cells were pelleted again by centrifugation and resuspended in PBS containing 1.25 μg/ml DAPI (Thermo Fisher cat no. D3571). Cells in PBS and DAPI were incubated for 5 min in the dark. Cells were then pelleted again and resuspended in 30 μl DAPI solution. Cells were imaged on a Nikon Eclipse TI2 automated fluorescent microscope using the NIS Elements Imaging Software (Version 5.21.02). Eighteen separate frames per well were captured using a Plan Apo λ 20x objective at 4×4 binning and 16-bit depth using brightfield microscopy and DAPI (excitation wavelength: 395 nm; 10ms exposure: 100% power) and green (excitation wavelength: 470 nm; 2ms exposure; 100% power) fluorescent channels.

### Data Analysis

Data analysis was conducted in R (v4.1.3). To reproducibly analyze the flow cytometry data, fcs files were read into R, transformed, and gated using the flowCore (2.6.0; [29]), flowViz (1.58.0; [30]), and ggcyto (1.22.0; [31]) packages. Gating-ML 2.0 [32] files documenting data transformations and gate coordinates were written with the flowUtils (1.58.0; [33]) package. For all samples, fluorescence intensity measures were transformed prior to gating with the “logicle” transformation [34], using the logicletGml2() function from flowCore and default parameters. To identify significant differences between various conditions, linear models were fit with the base lm() function and the Wald test was used to determine significance. MFI values were first Log_10_- transformed before linear modeling. For comparing the effect of NLS domains on GFP-fused enzyme localization, nuclei were segmented using the Nikon Elements software based on DAPI signal (nuclei were ∼9.6 μm in diameter). An annulus outside the nucleus was drawn to represent the cytoplasm (diameter of the outer ring of the cytoplasmic annuli were ∼11.3 μm) and the average (per pixel) GFP intensity within the nucleus and the cytoplasmic annulus (i.e. excluding the nucleus) was calculated separately. Cells with a total (nuclear and cytoplasmic) GFP intensity below 500000 pixel intensity were excluded as GFP-negative. All GFP-positive cells imaged in the well were quantified and the mean Log_2_-transformed ratio of nuclear-to-cytoplasmic GFP intensity was calculated for each of three biological replicates for each construct. Finally, significant differences in the Log_2_-ratio were determined by Wald tests in linear model fits of pairwise comparisons.

## Results

### Initial optimization of existing dCas9 transcriptional cascade system

As an initial step in evaluating CRISPR-driven gene regulatory networks, we sought to implement a previous dCas9 system [5]. To drive this cascade, dCas9 is directed in a defined order to a series of reporter plasmids containing synthetic promoters with tandem arrays of gRNA target sites. When the first reporter commences expression, the gRNA to activate the second reporter is encoded in the 3’ untranslated region (“UTR”) of the transcript for the first reporter gene and is cleaved at 28 nt recognition sites by the Csy4 nuclease (**Fig. 1A**, left side). After obtaining the available plasmids for this system from Addgene, we found that several optimizations were needed. First, we found that the plasmids were prone to leaky background expression when transfected at high concentration (i.e. fluorescent signal was detected from cells transfected with the downstream reporter but not the upstream activator), so we optimized the transfection conditions for HEK293T. Second, while the original publication reported results using EYFP, ECFP, and mKate2 as reporters, the plasmids the authors had made available on Addgene included PhiYFP, mCerulean, and dsRed2. Unfortunately, dsRed2 matures relatively slowly and has an intermediate green fluorescent stage [15,16] making it difficult to distinguish expression from the two stages of the cascade even with compensation. We initially tested several alternative red fluorescent proteins (FPs), including iRFP670 and mCherry (data not shown), but found them to exhibit various levels of leaky background expression even when expressed from the same promoter. Ultimately, mCherry had less background expression than iRFP670, but also exhibited some intrinsic fluorescence in the mCerulean spectrum that could not be properly compensated.

We proceeded with PhiYFP, mCerulean, and mCherry, but limited our analyses to using one fluorophore at a time. Future work would be required to identify or engineer a red FP without background expression or fluorescence signals in other spectra. Third, one of the constructs (“P2”) did not match the expected restriction digest pattern upon receipt and so we amplified the promoter and reporter gene region and inserted it into the backbone of the upstream plasmid in the cascade (“P1”). Finally, in addition to P1 and P2, we designed a new construct (“P3”), which uses a previously reported gRNA recognition site derived from an endogenous locus in the human genome [17] to construct a synthetic promoter consistent with P1 and P2.

### Development of a dCas12a transcriptional cascade system

Having validated the ability of the original system to drive expression of reporter genes in a cascade, we next sought to replace dCas9 with dCas12a as the driver of the cascade. To do so, we added dCas12a-[Activ], which utilizes a custom transactivating domain that combines components of p65 and HSF1 [17], to the initial construct in the cascade, and then converted the 28 nt Csy4 recognition sequences flanking the relevant guide RNAs with Cas12a DRs. Due to its intrinsic RNase activity, dCas12a can process the crRNAs to initiate the next reporter in the sequence, unlike dCas9, which requires Csy4 to process the gRNAs. While Cas9 recognizes an NGG protospacer adjacent motif (PAM) 3’ of its target site, Cas12a recognizes a TTTV PAM 5’ of its target site (the V indicates that the final base can be A, C, or G following the International Union of Pure and Applied Chemistry nomenclature [18]). A TTTC sequence in the synthetic tandem array promoters of the existing dCas9 cascade allowed us to target the same synthetic promoters with dCas12a targeting the opposite strand from dCas9 (**Fig. 1A**, right side).

We then tested the ability of both dCas9-VP64 and dCas12a-[Activ] to drive expression of reporter genes in the P1, P2, and P3 constructs (**Fig. 1, Fig. S1, Table S2**). Both dCas activator proteins were able to drive expression of all three reporters, with roughly similar levels of activity, both in terms of percent positive cells (%positive) and Log_10_-transformed median fluorescence intensity (MFI) for the fluorescent proteins among positive cells (**Fig. 1B-G**). Because dCas12a was tested across various amounts of transfected plasmid DNA, we fit a linear model through the %positive data at different amounts of input for dCas12a-[Activ] and calculated where that line intersected the mean %positive for dCa9-VP64 using 250 ng. For the three reporters, the dCas12-[Activ] linear model intersected the dCas9-VP64 data between 499 ng and 697 ng (697 ng, 499 ng, and 696 ng for P1, P2, and P3, respectively), and so we proceeded with comparing the two activators using 250 ng of dCas9-VP64 and 500 ng of dCas12a-[Activ]. We also noted that with all three reporters, the fluorescence range was broader and more uniform across cells when activated with dCas12a compared to dCas9 (**Fig. S2**).

To confirm that the observed reporter activities were not due to a nonspecific interaction between the activator proteins and the synthetic promoters (especially given the presence of PAM sequences in the promoters and the fact that our Cas12a crRNAs were identical for the first 9 bases for P1, P2, and P3), we further tested for cross-reactivity by transfecting each promoter along with its non-cognate guide RNAs (e.g. measuring fluorescence of cells co-transfected with P1 and guide RNA2 or guide RNA3). For SGRN, we found that P3 was mildly promiscuous (mean %positive = 5.7% for both crRNA1 and crRNA2), but that the other promoters had relatively high fidelity - mean %positive = 0.4% and 0.7% for P1 targeted by crRNA2 and crRNA3, respectively, and mean %positive = 2.1% and 2.0% for P2 targeted by crRNA1 and crRNA3, respectively (**Fig. S3, Table S3**). A highly similar pattern was observed for dCas9-VP64, suggesting that the promoter itself determines the degree of promiscuity more than the Cas enzyme or the mismatched guide RNA.

### Guide RNA length has different effects on transcriptional activation for dCas9 and dCas12a

Previous work from other groups has suggested that Cas enzymes have different relative activities of editing and transcriptional activation depending on the length of the guide used. Several studies have shown that guides need to be at least 17 nt for full editing efficiency [19–22], while Campa et al. reported that guides as short as 15 nt could still activate genes but were not capable of driving detectable editing [17]. Therefore, we decided to compare the relative activity of the Cas9-based cascade and SGRN when using guides of shorter lengths for the P2 promoter. We designed a series of constructs with 16 to 20 nt long guides and tested for the relative activation efficiency, again using both %positive cells and MFI as readouts (**Fig. 2, Table S4**). dCas9 had relatively consistent activity across the range of guide lengths in terms of %positive (linear modeling of %positive = guide length: slope = 0.93, p-value = 0.31). In contrast, dCas12a was much less active at 16 nt in terms of %positive, with increasing activity coinciding with increasing guide length (linear modeling of %positive = guide length: slope = 5.0, p-value = 1.1×10^-4^). We interpret this as being consistent with the reported higher specificity of dCas12a than dCas9 [14]. Interestingly, the effect of guide length on MFI was much less pronounced for dCas12a (linear modeling of Log_10_(MFI) = guide length: slope = 0.048, p-value = 0.01) than dCas9 (slope = 0.081, p-value = 7.5×10^-5^).

### Effect of different numbers of binding sites on activity of CRISPR cascades

For consistency with previous work, we had originally implemented these cascades with 6 or 7 guide RNA binding sites in the synthetic promoters (the number of binding sites was slightly different between P1 and P2, presumably due to mutations introduced during cloning of the original constructs). However, we were interested in determining what effect fewer binding sites might have on the behavior of these reporters. Therefore, we designed constructs with varying numbers of binding sites in the P2 promoter (from one to seven sites) and tested the activity of these constructs for both dCas9 and dCas12a (**Fig. 3, Table S5**). Because the Cas9 and Cas12a target sites were not perfectly interleaved in the original P2 construct (**Fig. S4**), we also included the original P2 construct as a control. Interestingly, in the negative controls that did not include a Cas activation plasmid, fewer binding sites corresponded to higher background reporter expression and higher apparent variability across replicates. On the other hand, in the presence of Cas-mediated activation, fewer binding sites in the synthetic promoter corresponded to slightly lower %positive and Log_10_(MFI) for both dCas9 and dCas12a. dCas12a appeared to have a flatter response to increasing numbers of binding sites, particularly in terms of Log_10_(MFI) (linear model slope = 0.04, p-value = 4.1×10^-5^) relative to dCas9 (linear model slope = 0.10, p-value = 6.8×10^-9^). Nonetheless, in the presence of a Cas activator, all synthetic promoters had expression levels that were clearly distinguishable from the background (linear model comparing %positive for 1xTS only vs 1xTS + dCas9 and Csy4 p-value = 3.5×10^-3^, 1xTS only vs 1xTS + dCas12a p-value = 4.4×10^-4^).

### Differences in promoters and activation domains do not fully explain differences in activity of dCas9 and dCas12a

While the two Cas systems were similar overall, there were several differences - including the amount of construct needed for equivalent activity, sensitivity to guide length, robustness to the number of target sites, and overall expression of reporter in positive cells. Given that [Activ] is reported to be a stronger activator than VP64, it was somewhat unexpected for dCas9-VP64 to match the activity of dCas12a-[Activ] with half the amount of plasmid included in transfections. Therefore, we evaluated the influence of individual components on activity. Because the two Cas proteins were expressed with different promoters (UbCp and EF1α for dCas9 and dCas12a, respectively) and utilized different transcriptional activation domains (VP64 and [Activ] for dCas9 and dCas12a, respectively), we first designed an experiment to test the effect each element had independently on the activity of our dCas12a system. We compared all four promoter and activator combinations for dCas12a targeting P2 (**Fig. 4, Table S6**). Fitting linear models to both the %positive and Log_10_(MFI) dCas12a results (including terms for promoter and transcriptional activation domain), we found that expressing dCas12a from the UbCp promoter indeed resulted in significantly lower activity (%positive p-value = 2.21×10^-5^ and Log_10_(MFI) p-value = 1.24×10^-7^) compared to the EF1α promoter, consistent with previous reports [23]. The models indicated that relative to the EF1α promoter, UbCp was responsible for a reduction in %positive of 13.8 and a reduction in MFI of 1758.0. On the other hand, while the [Activ] and VP64 transcriptional activation domains were also significantly different (%positive p-value = 0.027 and Log_10_(MFI) p-value = 4.71×10^-5^), with VP64 associated with an increase in %positive of 4.6 and an increase in MFI of 1806.6, contrary to previous reports [10]. We also tested the relative activity of [Activ] and VP64 with the P1 and P3 reporters (**Fig. S5, Table S7**). For P1, VP64 was significantly less active in terms of Log_10_(MFI) (p-value = 4.2×10^-4^), with a slight reduction in MFI of 552.9, but the two activators were not significantly different in terms of %positive (p-value = 0.34, **Fig. S5B-C**). For P3, the two transcriptional activators were not significantly different for either %positive (p-value = 0.34) or Log_10_(MFI) (p-value = 0.93, **Fig. S5D-E**).

Because the synthetic promoters use arrays of target sites with relatively few bases between the targets for spacing, we considered whether differences in size (and potential steric hindrance at the promoters) of the two activators might affect their overall activity (given that dCas12a-[Activ] activator is approximately 14.7 kDa bigger than the dCas9-VP64 activator - 176.3 kDa vs 161.6 kDa, respectively). We therefore compared the efficiency of the two activators when targeting either a promoter with one (1xTS) target site or a promoter with seven (7xTS) target sites (**Fig. S6, Table S8**). Fitting linear models to test the effect of the transcriptional activators on the 1xTS promoters and 7xTS promoters separately, we did observe significant differences in Log_10_(MFI) that could be consistent with steric hindrance. We found VP64 less active on the 1xTS promoter (p-value = 0.017) and more active on the 7xTS promoter (p-value = 0.013). However, the effect sizes were relatively small and no significant differences were observed in the %positive values, indicating that a steric hindrance effect alone was insufficient to explain the patterns we observed in transcriptional activity between the two activators.

### Differences in nuclear localization explain major differences in activity of transcriptional activation systems

While differences in activation domains and promoters did not fully explain differences in the activity of the dCas9 and dCas12a systems, others in the literature have noted that the editing efficiency of Cas12a is greatly improved by more localization of the enzyme to the nucleus [24,25]. Notably, the original dCas12a-[Activ] construct contained a single NLS [17], compared to three NLSs present in the dCas9-VP64 construct. To evaluate whether localization might explain differences, we first sought to visualize the effect that incorporating different NLS peptide sequences might have on cellular localization. We first fused GFP to various dCas activating enzymes - including dCas9-VP64, dCas12a-[Activ] (1xNLS), dCas12a-3xNLS-[Activ] (using the same NLS as dCas9), and dCas12a-3xNLS-VP64 - and visualized their localization. dCas12a- [Activ] (1xNLS) was significantly less localized to the nucleus than any of the other GFP constructs (**Fig. 5A, Fig. S7, Table S9**). dCas9-VP64 and dCas12a-3xNLS-VP64 achieved the best nuclear localization. To test the effect of differential localization on transcriptional activation, we tested several dCas constructs (without GFP fusions), including dCas9-VP64 (3xNLS), dCas12a-[Activ] with 1x or 3x NLS, and dCas12a-VP64 with 1x or 3x NLS. All constructs were transfected in triplicate and evaluated for %positive and MFI as in the previous experiments (**Fig. 5B-D, Table S10**). Fitting a linear model to the dCas12a results (including terms for both the activation domain and the NLS copy number) identified a significant increase in %positive rates for 3xNLS relative to 1xNLS (p-value = 2.4×10^-7^, increase in %positive of 22.1) and Log_10_(MFI) (p-value = 2.2×10^-10^, increase in MFI of 6304.5). In addition, [Activ] drove a subtle but significantly higher Log_10_(MFI) than VP64 (p-value = 4.3×10^-3^, increase in MFI of 215.9). In parallel with this experiment, we also repeated the titration of the dCas12a constructs (testing three input masses for [Activ]-1xNLS, [Activ]-3xNLS, and VP64-3xNLS, **Fig. S8, Table S10**). While dCas9 had slightly lower activity this time relative to [Activ]-1xNLS, both of the 3xNLS constructs greatly exceeded dCas9-VP64, even at the same input (250 ng). Interestingly, with improved localization of dCas12a using the 3xNLS, VP64 produced significantly higher %positive values than [Activ] (p-value = 0.025), while [Activ] drove significantly higher Log_10_(MFI) than VP64 (p-value = 5.2×10^-5^).

### Nuclear localization improves expression of three-level cascade

Given the improved performance of dCas12a with improved nuclear localization, we decided to investigate how this signal might propagate through a higher-order cascade. Therefore, we designed multiple three-level cascades utilizing either dCas9 or dCas12a. For each system, the first level encodes a transcript that includes a guide RNA (gRNA1/crRNA1). For dCas9, this transcript does not have a protein-coding gene. For dCas12a, this transcript also expresses dCas12a itself. These guide RNAs then target the second level promoter (P1), and the second level transcript includes a guide RNA (gRNA2/crRNA2) targeting the third level promoter (P2). The transcript downstream of P2 encodes mCerulean, which provides a readout for completion of the three-level cascade (**Fig. 5E-G, Table S11**). In this experiment, we tested dCas12a fused to both activation domains, dCas12a-VP64 and dCas12a-[Activ] (both with a 3xNLS). We found that the dCas12a-[Activ] construct yielded significantly greater %positive and Log_10_(MFI) values than dCas9 (%positive p-value = 0.010; Log_10_(MFI) p-value = 4.8×10^-4^) and dCas12a-VP64 (%positive p-value = 0.021; Log_10_(MFI) p-value = 6.1×10^-4^). Overall, this suggests that while we do not see the same degree of increased activity for [Activ] on primary targets that others have reported, [Activ] is able to drive higher reporter expression for three-level cascade systems.

## Discussion

In this work, we developed a synthetic transcriptional cascade system that we call SGRN. The system depends on dCas12a to drive expression of target genes in a predetermined order. Our system builds on previous work by streamlining an existing dCas9 approach [5]. While dCas9 initially drove higher expression of target genes, testing several parameters led to the discovery that nuclear localization was a critical component for the efficiency of the system. In fact, we found that different transactivating domains could affect nuclear localization, confounding the interpretation of head-to-head comparisons of these domains. Importantly, with better localization, dCas12a activity far exceeded the activity we observed for dCas9. In addition, our dCas12a method benefits from several advantages. First, we optimized both systems for minimal background expression, established rigorous flow cytometry parameters for evaluating the effect of the system, and empirically determined that single-target synthetic promoters can be used to rapidly generate any number of suitable guide RNA sequences. Although the system is more robust if subsequently implemented with a target site array. Second, the implementation of dCas12a renders the inclusion of Csy4 unnecessary, and the shorter DR and crRNA sequences required for dCas12a function, relative to 28 nt and tracrRNAs in the dCas9 system, allow for more flexibility and greater efficiency in transfection and transduction settings. Finally, by expanding the repertoire of available CRISPR-based transcriptional regulatory systems, we have opened the possibility of more complicated regulatory systems that take advantage of dCas9 and dCas12a simultaneously, potentially like other work that has developed synthetic metastable states [26].

There are several caveats worth keeping in mind while evaluating these results. First, all experiments were carried out in the HEK293T cell line. We have not determined whether these observations will hold true across different cell types. Second, we used two measures of FPs to track the efficiency of the system - the percentage of cells detected as positive for a given FP and the median fluorescence intensity for positive cells. While both capture important phenomena, neither is a perfect descriptor of the data. In fact, we found the distribution of intensities for positive cells, and the relationship between fluorescence intensity and side scatter to reliably separate the two expression systems for reasons that are not totally clear. Furthermore, we chose protein measurements as proxies for function as we ultimately hope to use this system to drive expression of protein-coding genes. However, transcript levels may also help to understand the kinetics and dynamics of the system, especially for the guide RNA-containing transcripts. Finally, it is worth pointing out that all these experiments were conducted with transient transfections, in which many copies of each plasmid would initially enter the cell and then be diluted by cell doubling. The constraints of a single integration system (like lentivirus) might lead to different optimization decisions.

In the future, we hope to further modify this system to allow for integration in the genome, to make it inducible and selectable, to allow tuning of both the timing of switches between cascade levels and expression levels of individual proteins in the cascade, and ultimately to implement this workflow to generate genuine artificial cascades with biologically relevant proteins. We hope this work serves as a resource for the community and a springboard for further investigations into the utility of CRISPR/Cas systems for synthetic biology applications.

## Data Availability

Flow cytometry data will be made available from FlowRepository (http://flowrepository.org). Imaging data will be made available from OSF (https://osf.io).

Analysis code will be made available from OSF (https://osf.io) and Github.

A set of plasmids enabling cloning of arbitrary target sites and/or guide RNAs to facilitate building of custom cascades (see Methods for details) will be made available from Addgene (Addgene IDs 202041-202049). Other constructs described in this manuscript may be made available through Addgene as well, depending on requests from the community.

## Funding

This work was supported by the National Institutes of Health [NIGMS R35GM137896 to D.A.C. and NIGMS R35GM147128 to C.A.T.]; the Flow Cytometry Shared Resource is jointly supported by RII (Research Innovation & Impact) and UA Cancer Center [NCI P30CA023074].

## Acknowledgements

We would like to thank C. Romanoski for critical reading of the manuscript, kindly sharing reagents, and insightful advice. We would like to G. Mouneimne for kindly sharing reagents. We would like to thank the Cusanovich and Thorne labs for support and advice.

## Supplementary Figures

**Figure S1.**
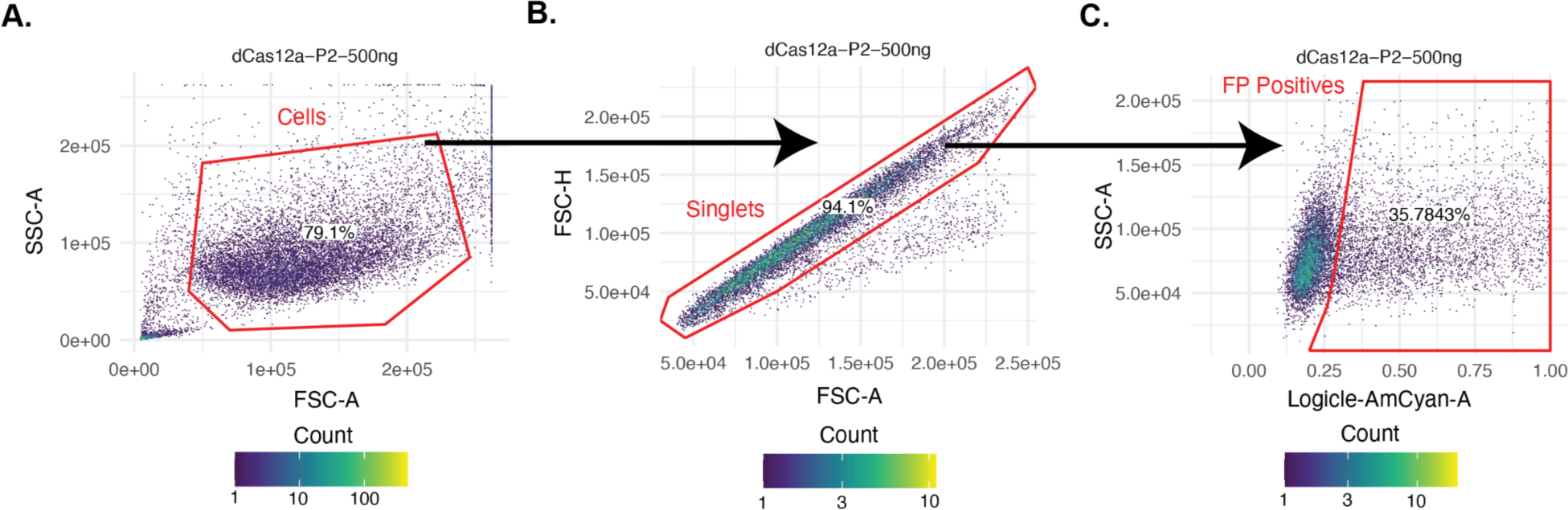
Example workflow for flow cytometry analysis. (**A**) Viable cells are gated based on forward scatter (FSC) and side scatter (SSC). (**B**) Singlets are gated based on area and height of forward scatter. (**C**) %positive and MFI are determined based on distributions in the appropriate fluorescent channel. Values shown here are from dCas12a-P2-500ng for replicate set 1 of the experiment conducted for Figure 1.

**Figure S2.**
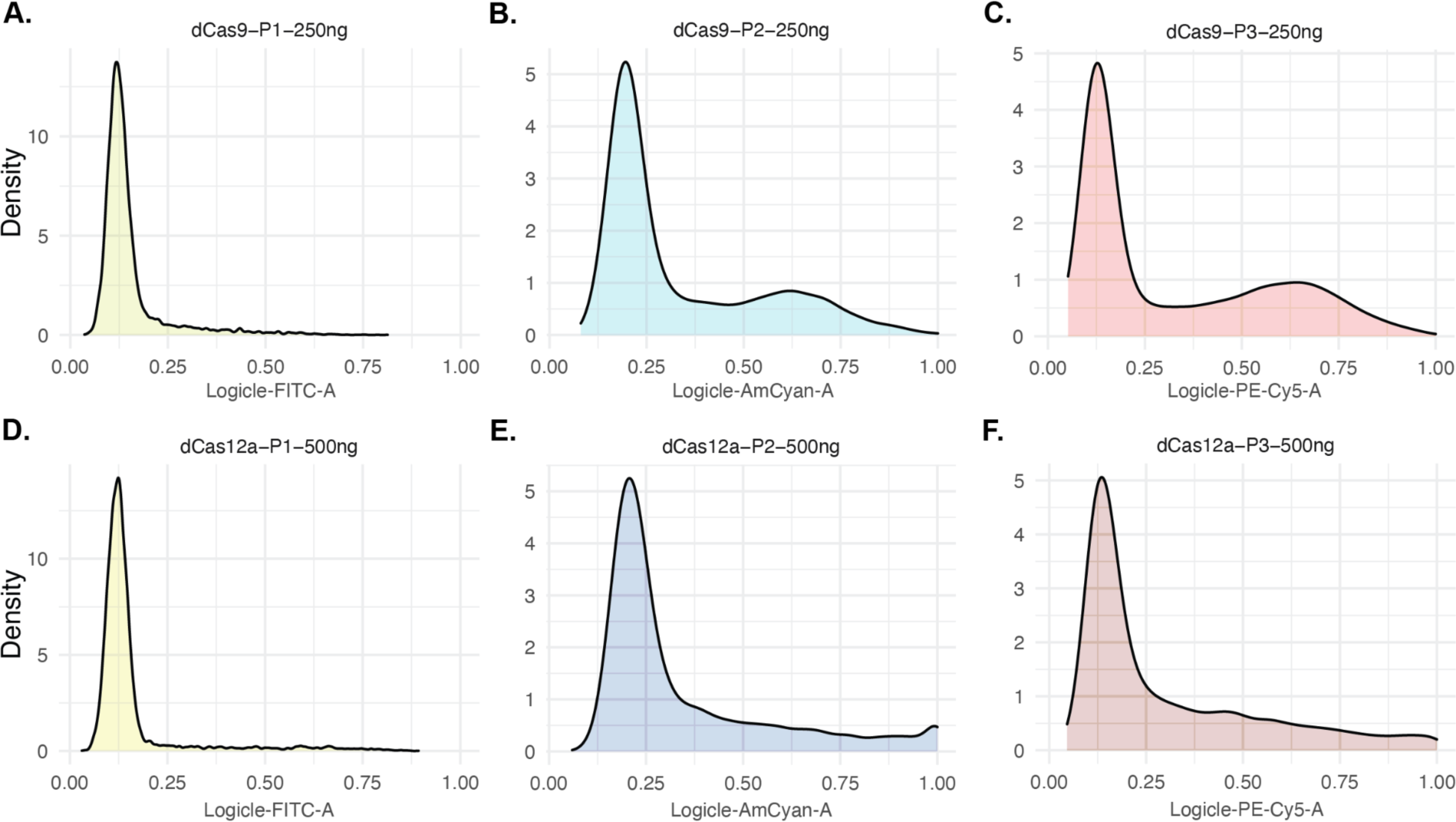
Distribution of fluorescence intensities for dCas9 and dCas12a targeting three different reporter constructs. (**A-C**) Fluorescence intensity density plot for dCas9-VP64 and Csy4, plus the indicated target reporter construct. (**D-F**) Fluorescence intensity density plot for dCas12a-[Activ] plus the indicated target reporter construct. (**A,D**) Fluorescence for P1 expressing PhiYFP. (**B,E**) Fluorescence for P2 expressing mCerulean. (**C,F**) Fluorescence for P3 expressing mCherry. Fluorescence values were transformed with the logicle algorithm. Values shown here are from dCas9 (250ng) and dCas12a (500ng) for replicate set 3 of the experiment conducted for Figure 1.

**Figure S3.**
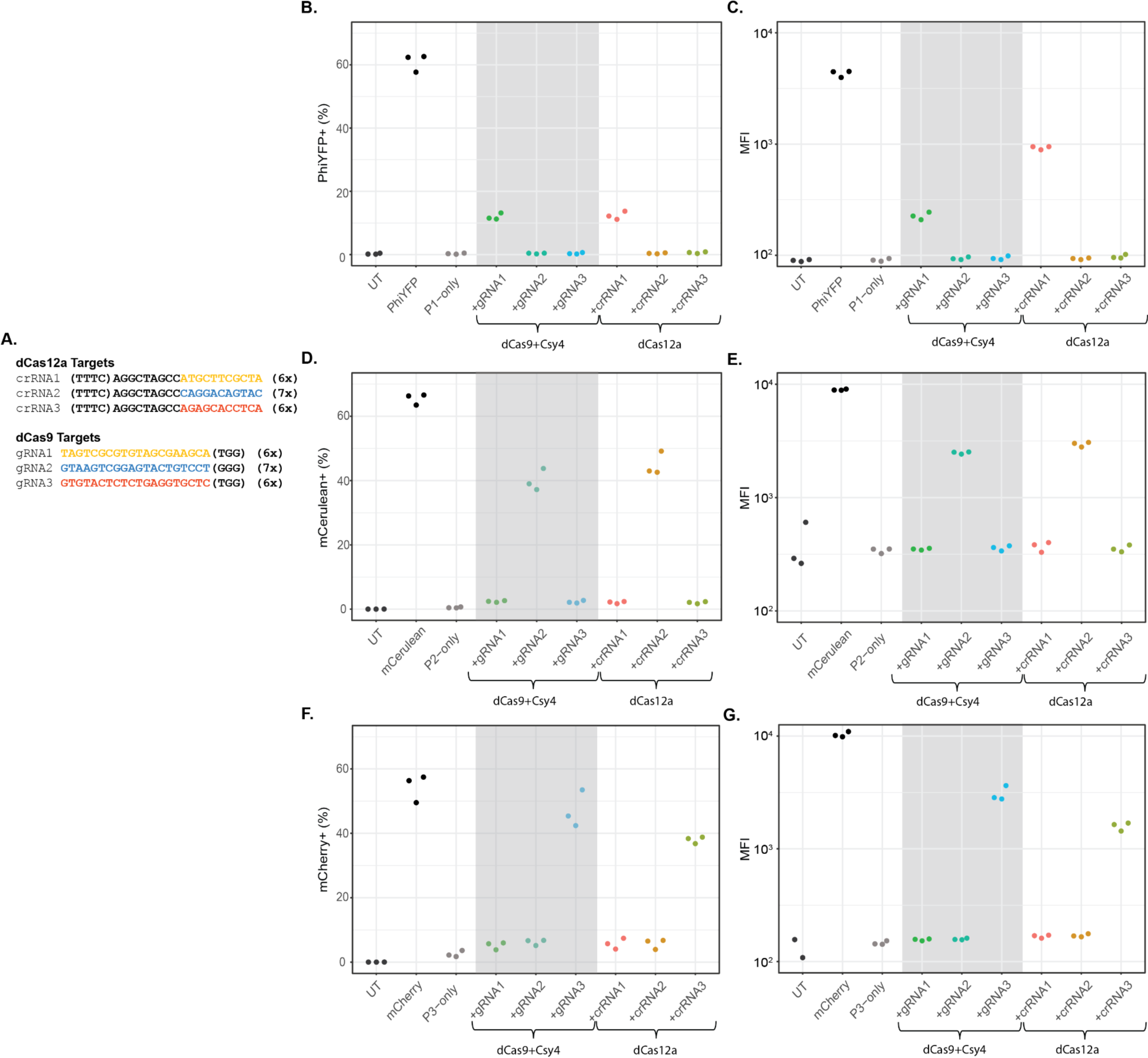
Testing cross-reactivity of guide RNAs. (**A**) Sequence of target sites. Colored bases indicate sequences that are unique to that target site. Black bases indicate sequences that are common to all target sites. Numbers in parentheses (e.g. 6x) indicate the number of replicates in the tandem arrays of these sequences that make up synthetic promoters P1, P2, and P3. (**B,C**) %positive and MFI (on Log_10_ scale) values for P1 “targeted” by the three different guide RNAs (crRNA1/gRNA1, crRNA2/gRNA2, and crRNA3/gRNA3). (**D,E**) %positive and MFI (on Log_10_ scale) values for P2 “targeted” by the three different guide RNAs. (**F,G**) %positive and MFI (on Log_10_ scale) values for P3 “targeted” by the three different guide RNAs. Note that one replicate for the UT sample in the P3 experiment is not plotted because there were no positive cells. “UT” denotes “untransfected”, “PhiYFP”, “mCerulean”, and “mCherry” denote constitutively expressed reporters that use the CMV promoter. To assess background transcription from the reporter genes, the downstream target plasmid was transfected without the upstream activator for each reporter - denoted as “P1-only”, “P2-only”, or “P3-only”, respectively.

**Figure S4.**
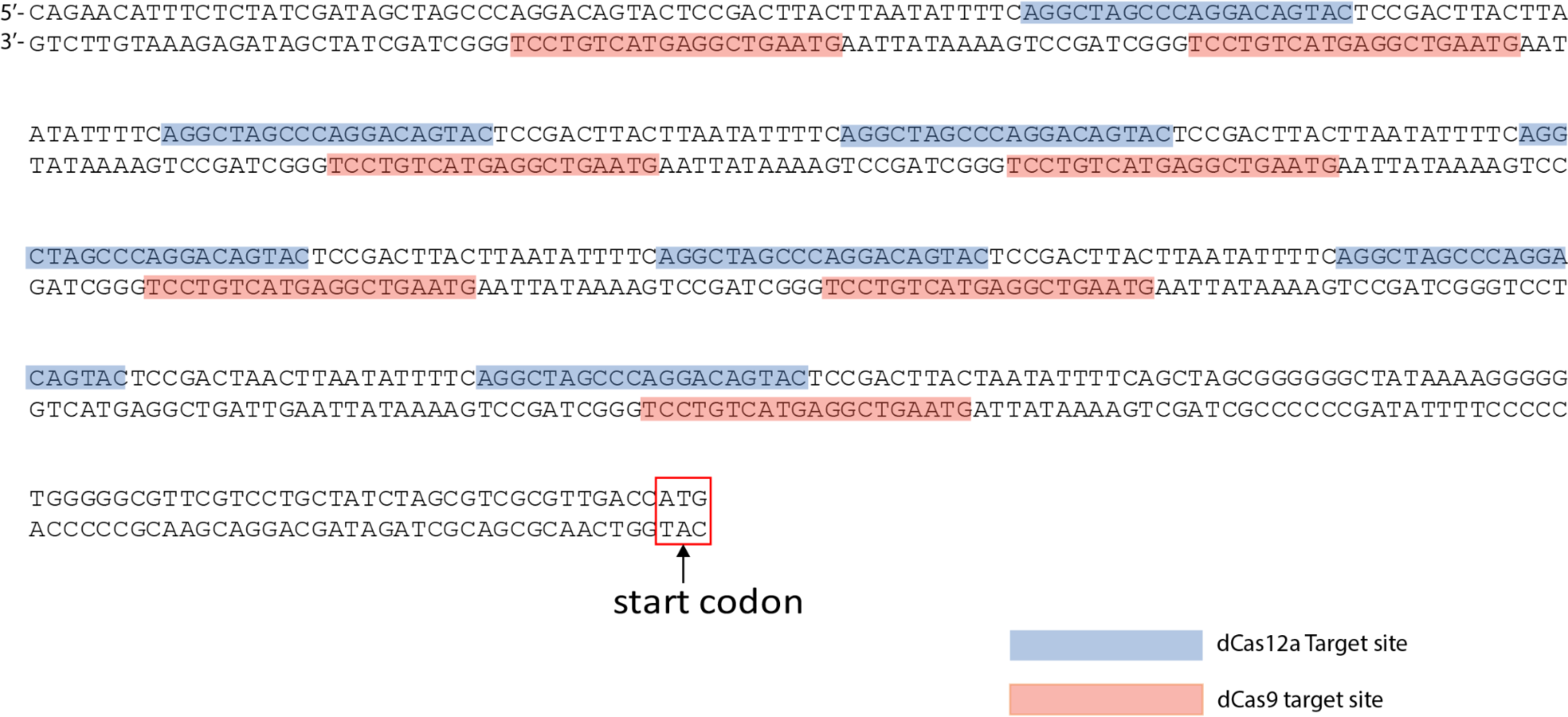
Target site locations of dCas9 and dCas12a binding in P2 promoter. Sequence of a region of the P2 construct encompassing the first target site for dCas9 to the start codon of mCerulean. dCas9 target sites (on the minus strand) are colored red. dCas12a target sites (on the plus strand) are colored blue. The PAMs are not highlighted, though the dCas9 PAM is 3’ of the target site and the dCas12a PAM 5’ of the target site.

**Figure S5.**
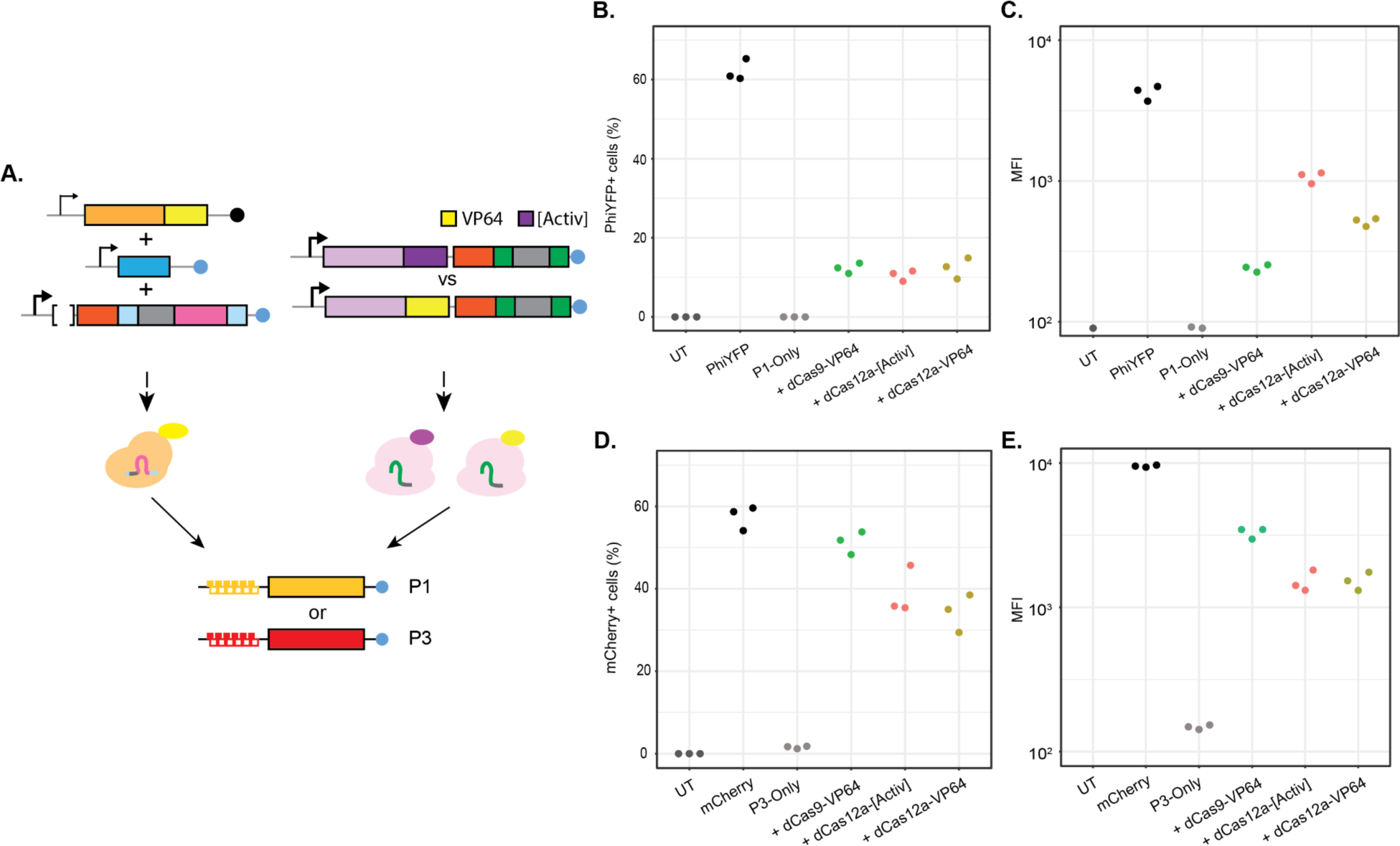
Effect of VP64 and [Activ] on different promoters. (**A**) Schematic of study design. dCas12a fused to either VP64 or [Activ] was compared to dCas9-VP64 for their ability to activate expression of P1 or P3. (**B,C**) %positive and MFI (on Log_10_ scale) values for P1 targeted by the three different activators. (**D,E**) %positive and MFI (on Log_10_ scale) values for P3 targeted by the three different activators. Note that several replicates in both experiments are not plotted because there were no positive cells for that condition. “UT” denotes “untransfected”, “PhiYFP” and “mCherry” denote constitutively expressed reporters that use the CMV promoter. To assess background transcription from the reporter genes, the downstream target plasmid was transfected without the upstream activator for each reporter - denoted as “P1-only” or “P3- only”, respectively.

**Figure S6.**
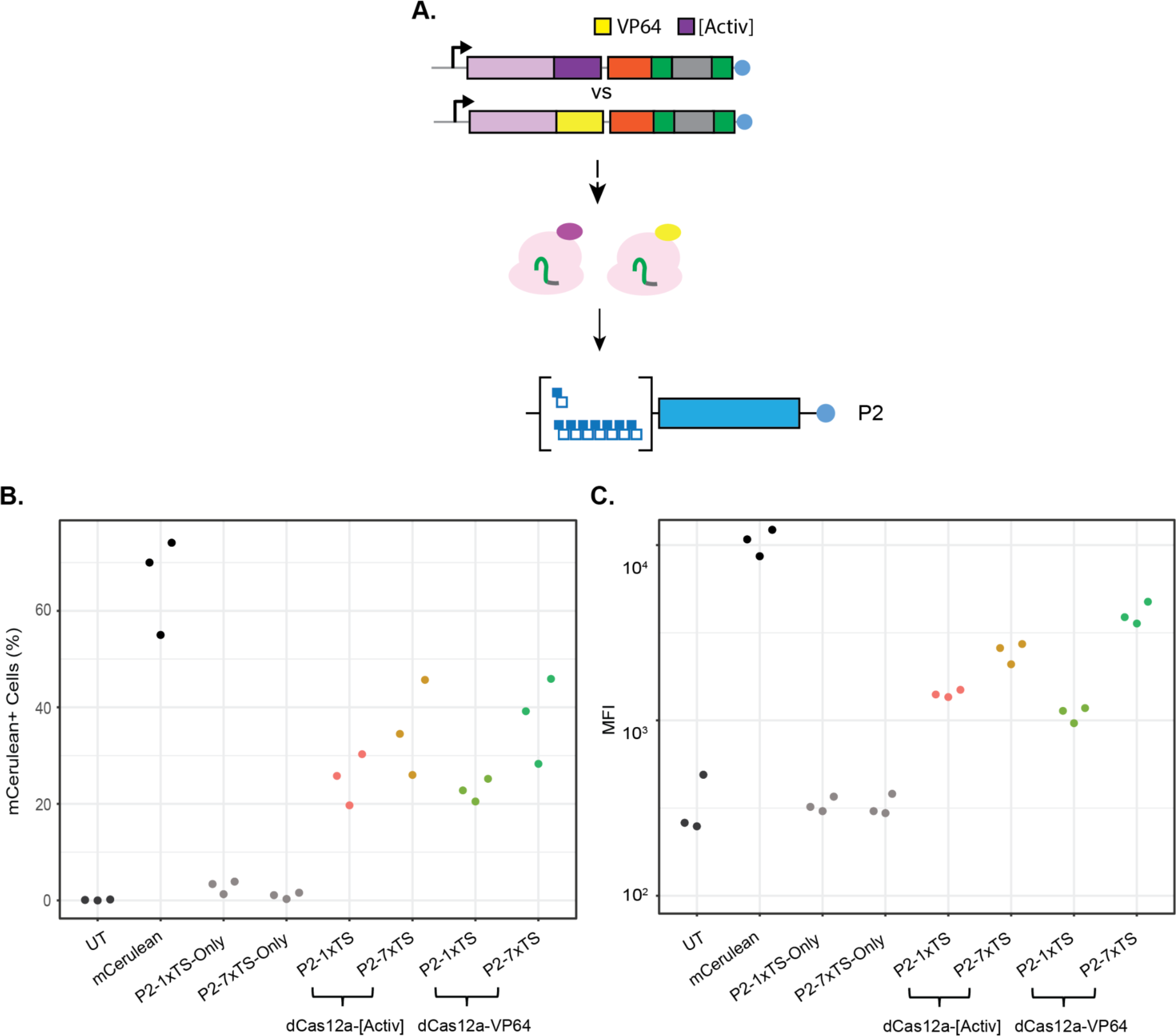
Evaluating effect of steric hindrance on activation domains. (**A**) Schematic of study design. dCas12a fused to either VP64 or [Activ] was co-transfected with versions of the P2 promoter that contained either one or seven binding sites for the Cas enzymes. (**B,C**) %positive and MFI (on Log_10_ scale) values for both P2 targets activated by the two different activators. “UT” denotes “untransfected”, “mCerulean” denotes a constitutively expressed reporter that uses the CMV promoter. To assess background transcription from the reporter gene, the downstream target plasmid was transfected without the upstream activator for each reporter - denoted as “P2-1xTS-only” or “P2-7xTS-only”, respectively.

**Figure S7.**
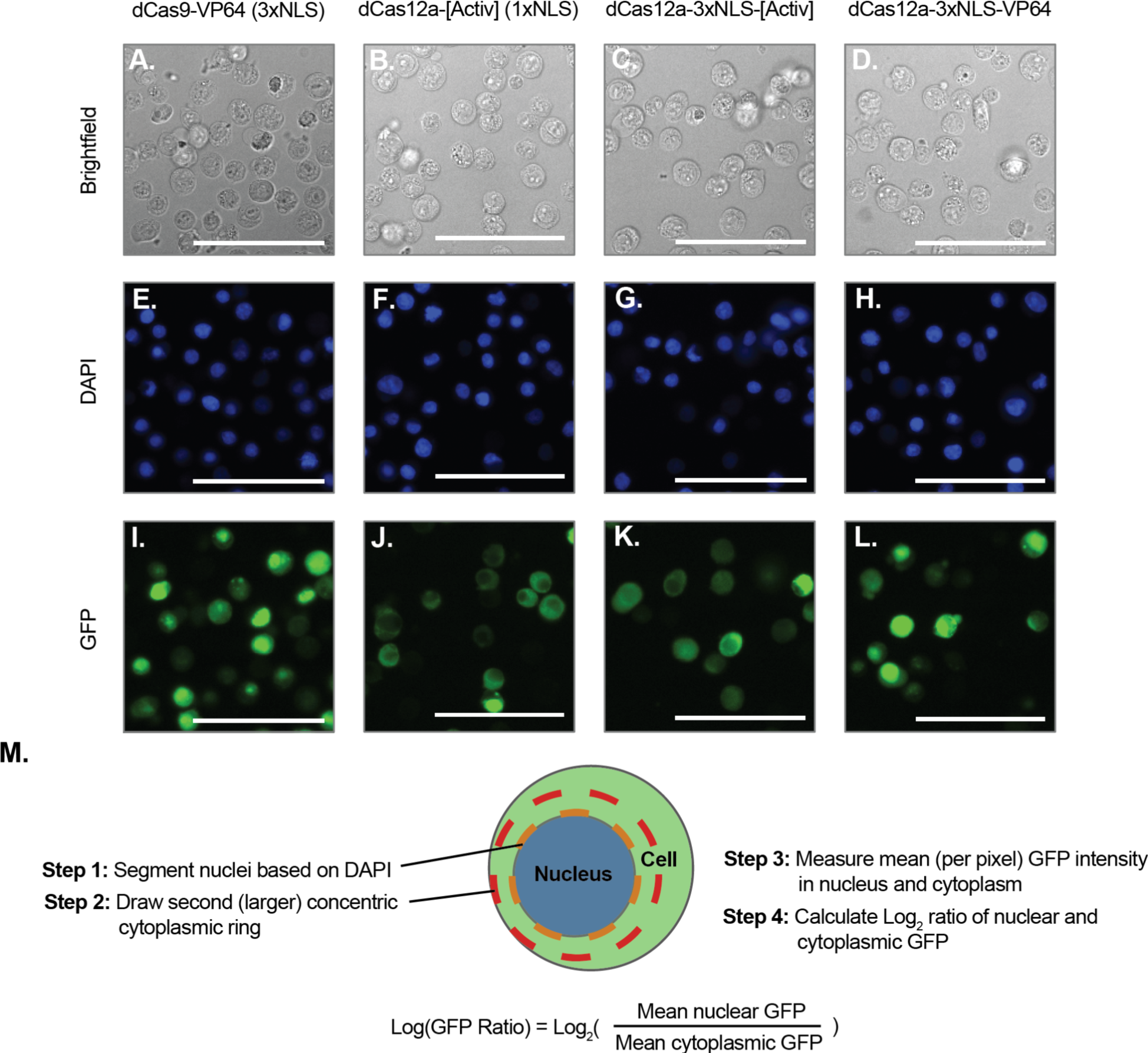
Nuclear localization of GFP-tagged dCas enzymes with different NLS. Cells were transfected with a plasmid expressing GFP-tagged versions of dCas9-VP64, dCas12a-[Activ], dCas12a- 3xNLS-[Activ], or dCas12a-3xNLS-VP64. 24 hours later cells were trypsinized, fixed, and stained with DAPI. (**A-D**) Brightfield images of cells transfected with each of the four constructs. (**E-H**) DAPI images of cells transfected with each of the four constructs. (**I-L**) GFP images of cells transfected with each of the four constructs. Scale bars indicate 100 μm for all images. (**M**) Schematic of GFP localization quantification analysis.

**Figure S8.**
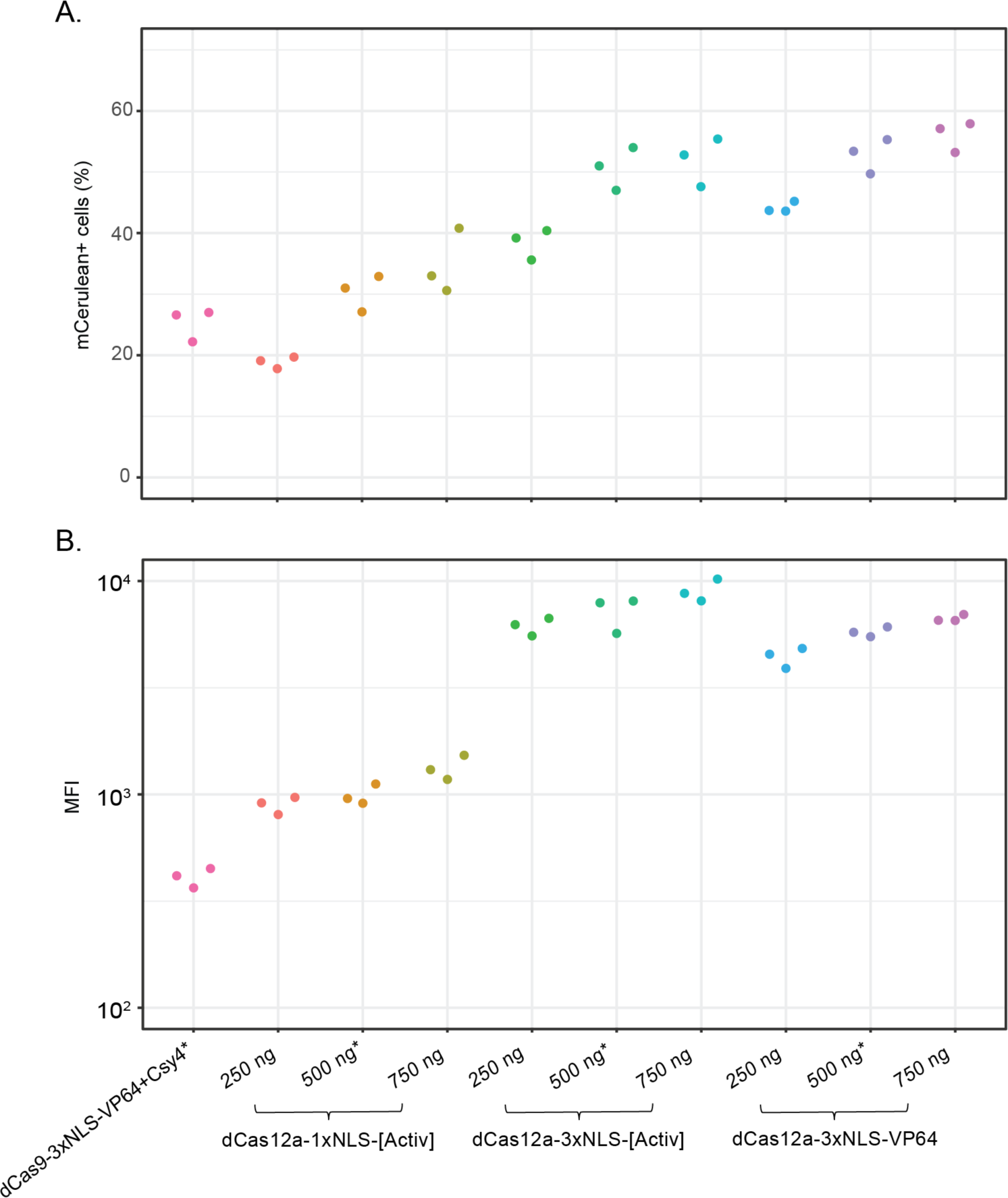
Titrating activity of 3xNLS-based dCas12a activators. (**A**) mCerulean %positive measurements for various constructs. The first column shows replicate values for dCas9-VP64 + Csy4 (the original Cas-based transcriptional cascade system). The remaining columns show the activity for various dCas12a-based activators tested across three different input amounts (250, 500, or 750 ng). Note that the values for dCas9 and the 500 ng inputs are the same data displayed in Figure 5, as both these experiments were run simultaneously. (**B**) MFI values (on a Log_10_ scale) for the same constructs (y-axis is displayed on a Log_10_ scale).

## Supplementary Tables

**Table S1.**
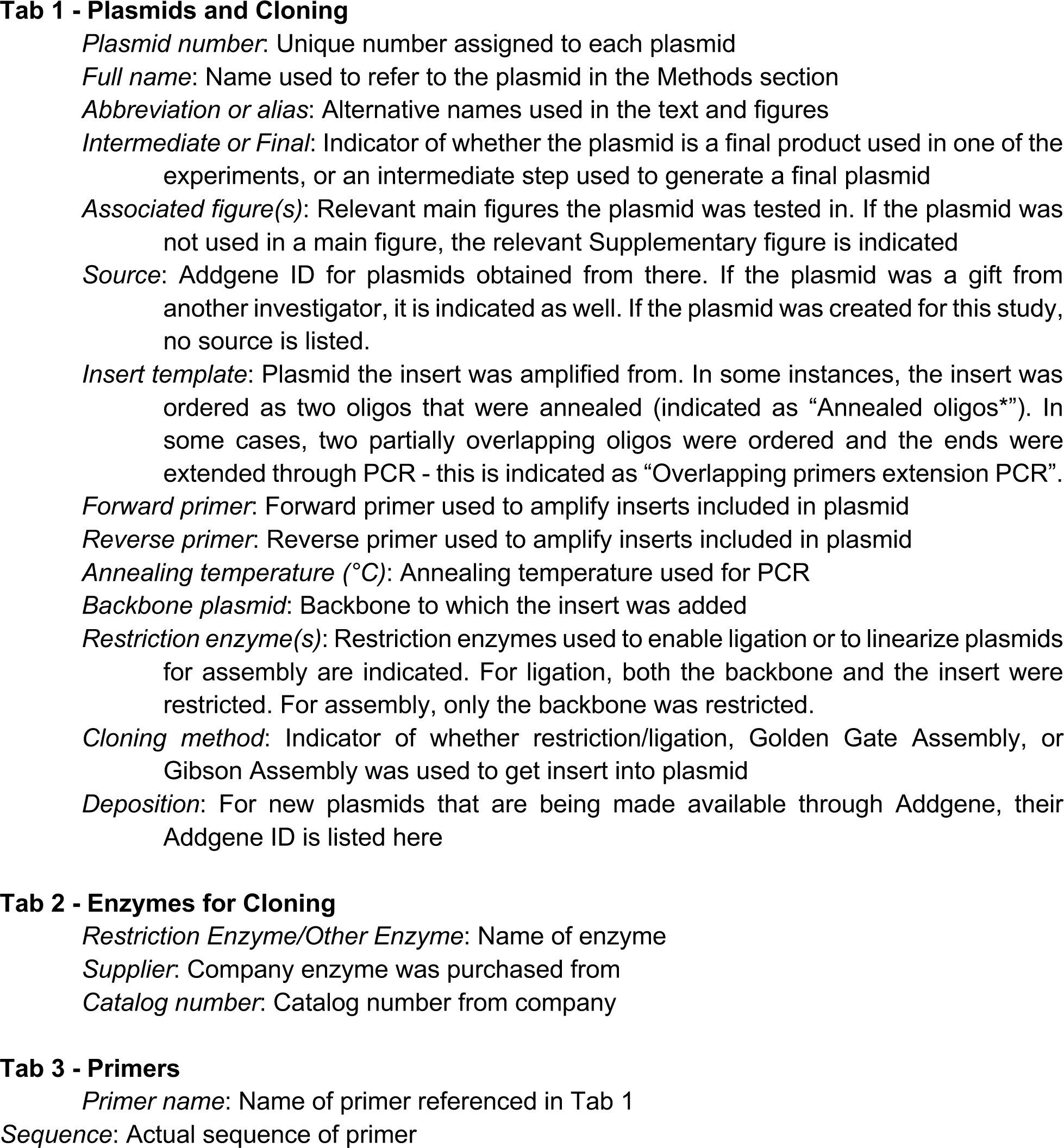
Details of cloning. This table is an Excel spreadsheet with three tabs. The columns of each tab are as follows:

**Table S2.**
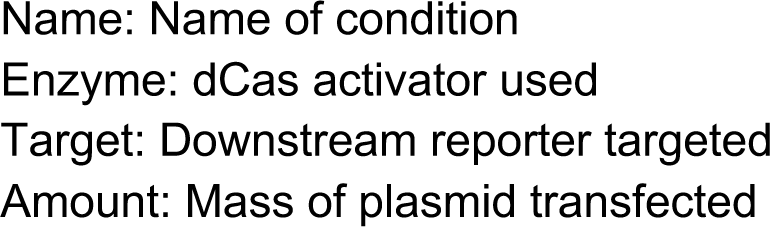
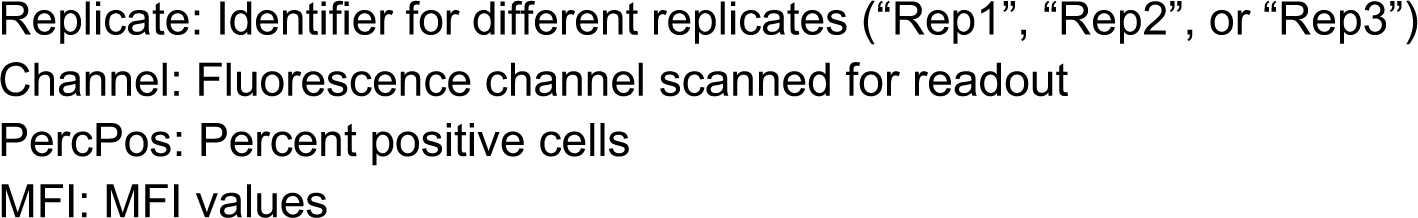
Initial comparison of dCas9 and dCas12a (relevant to Figure 1)

**Table S3.**
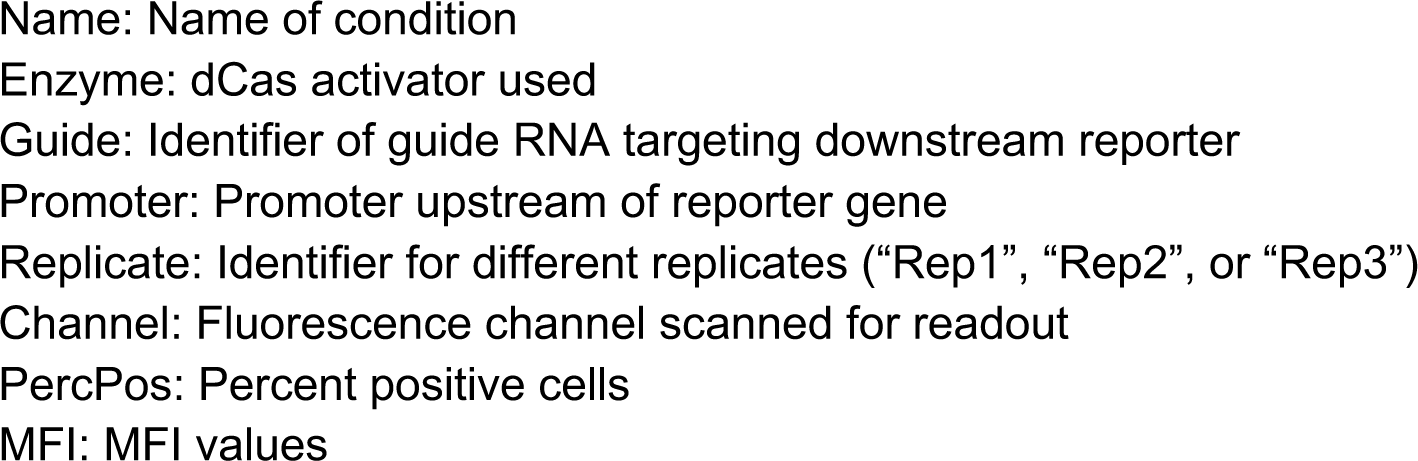
Tests of cross-reactivity of dCas9 and dCas12a (relevant to Figure S3)

**Table S4.**
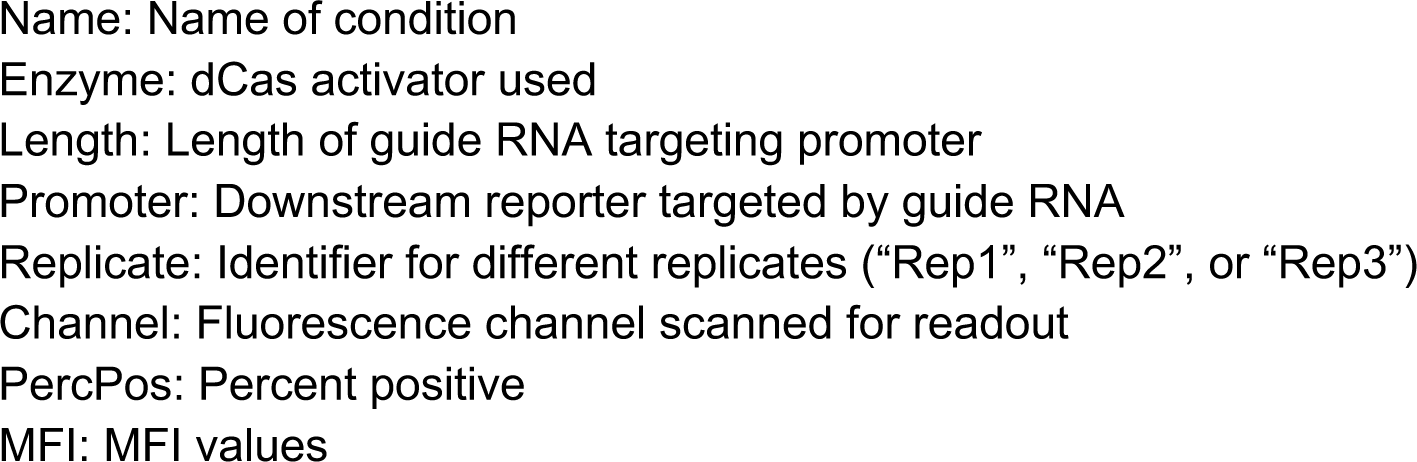
Effect of guide RNA length (relevant to Figure 2)

**Table S5.**
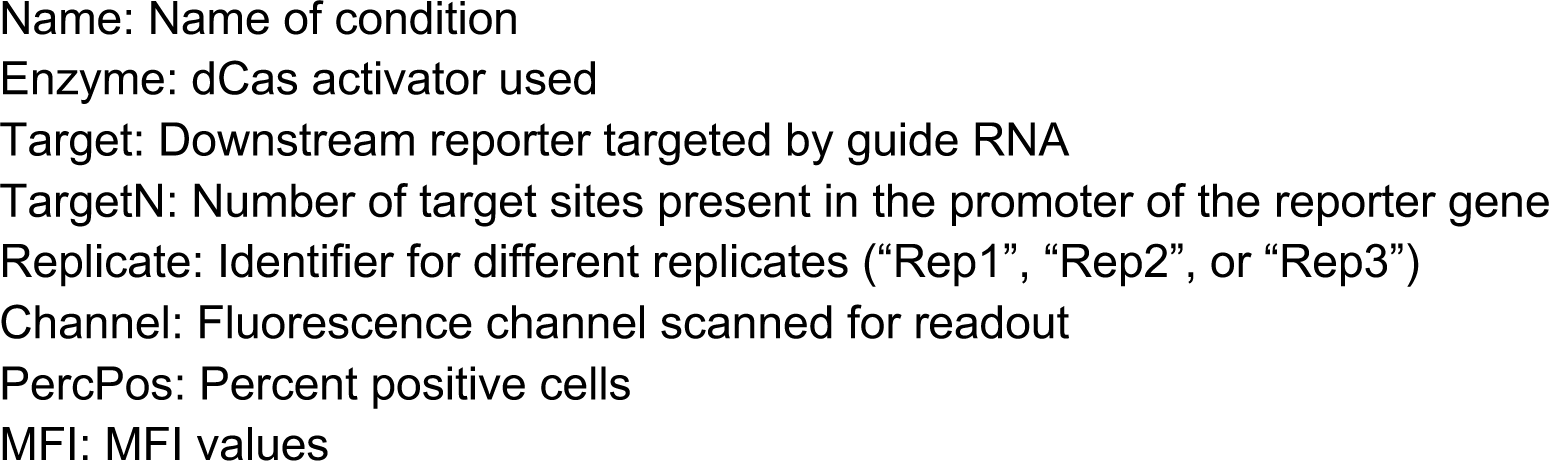
Effect of number of target sites (relevant to Figure 3)

**Table S6.**
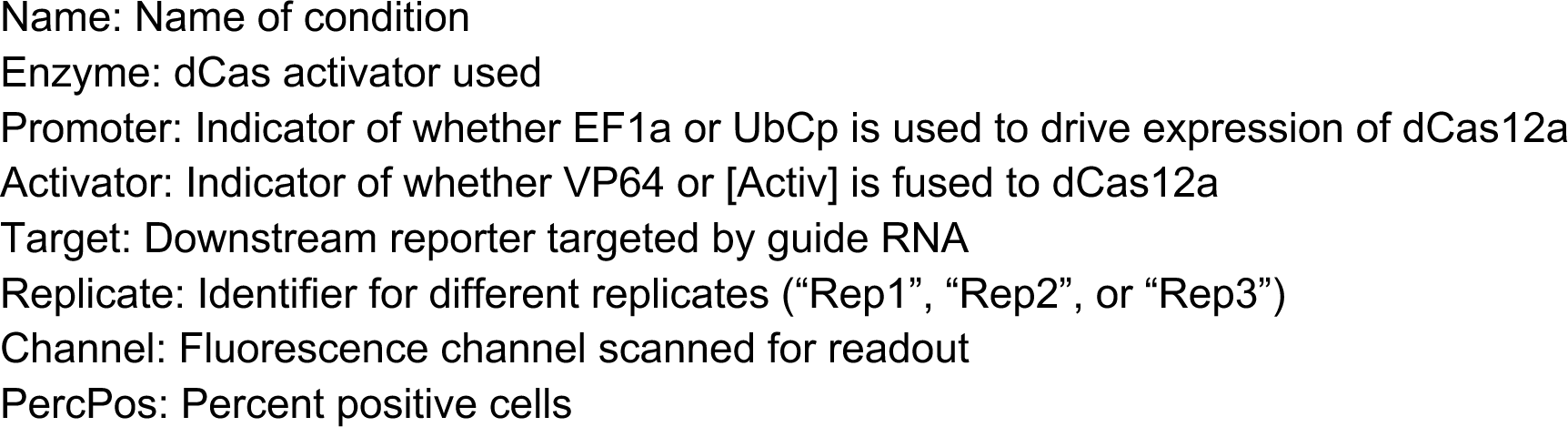
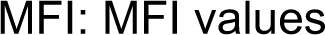
- Effect of domains on dCas12a activity (relevant to Figure 4)

**Table S7.**
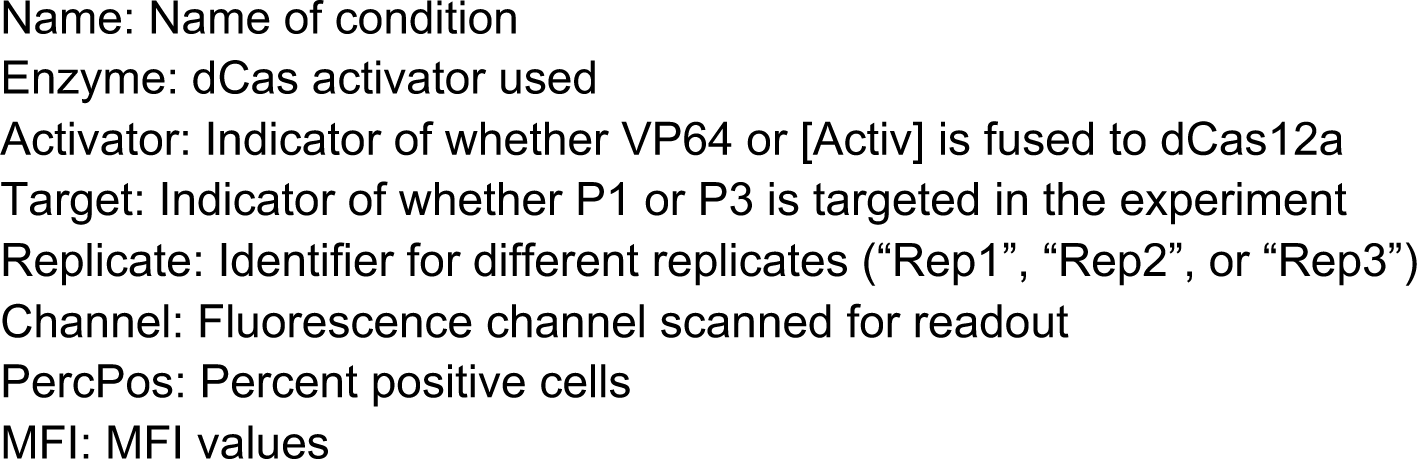
Effect of transactivation domain on dCas12a activity for P1 and P3 (relevant to Figure S5)

**Table S8.**
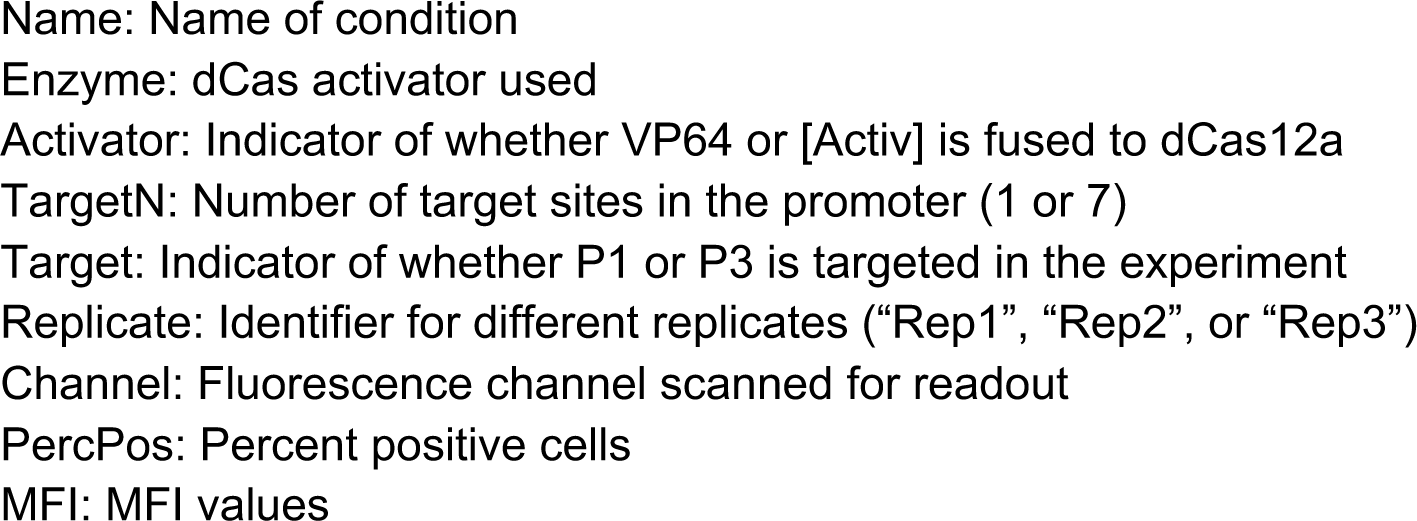
- Effect of steric hindrance on dCas12a activity for different transactivation domains (relevant to Figure S6)

**Table S9.**
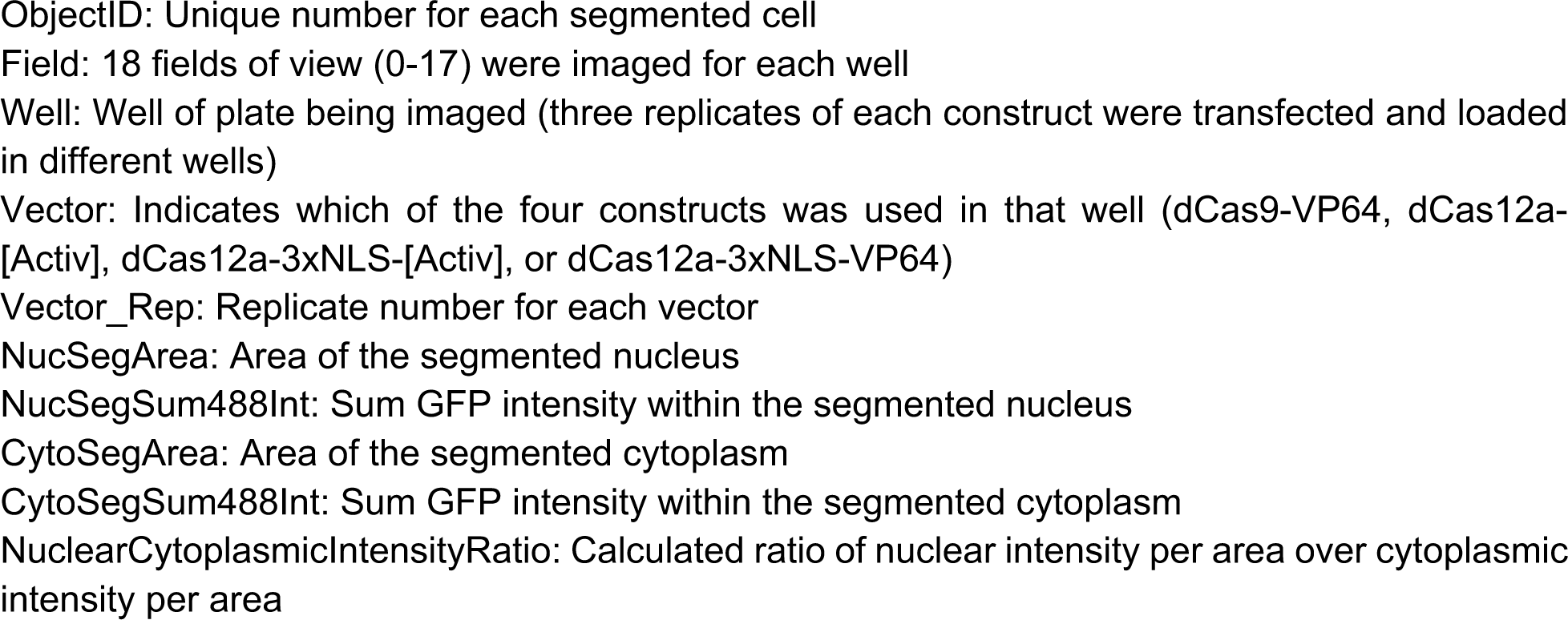
Single-cell Imaging data (relevant to Figure 5)

**Table S10.**
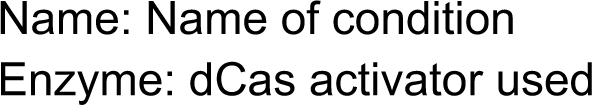
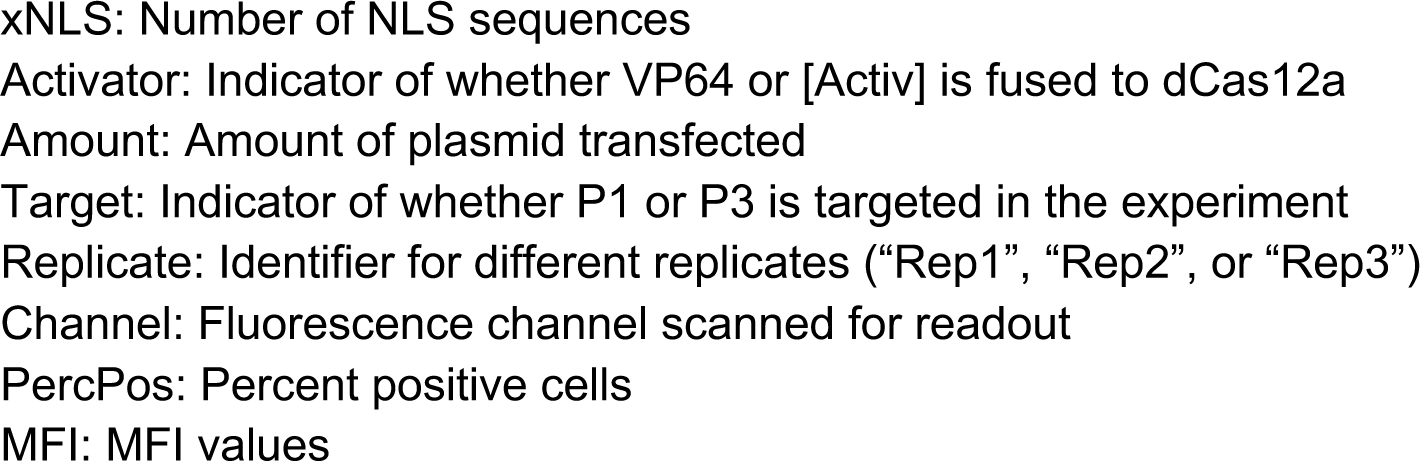
Effect of NLS domains on dCas12a activity (relevant to Figures 5, S8)

**Table S11.**
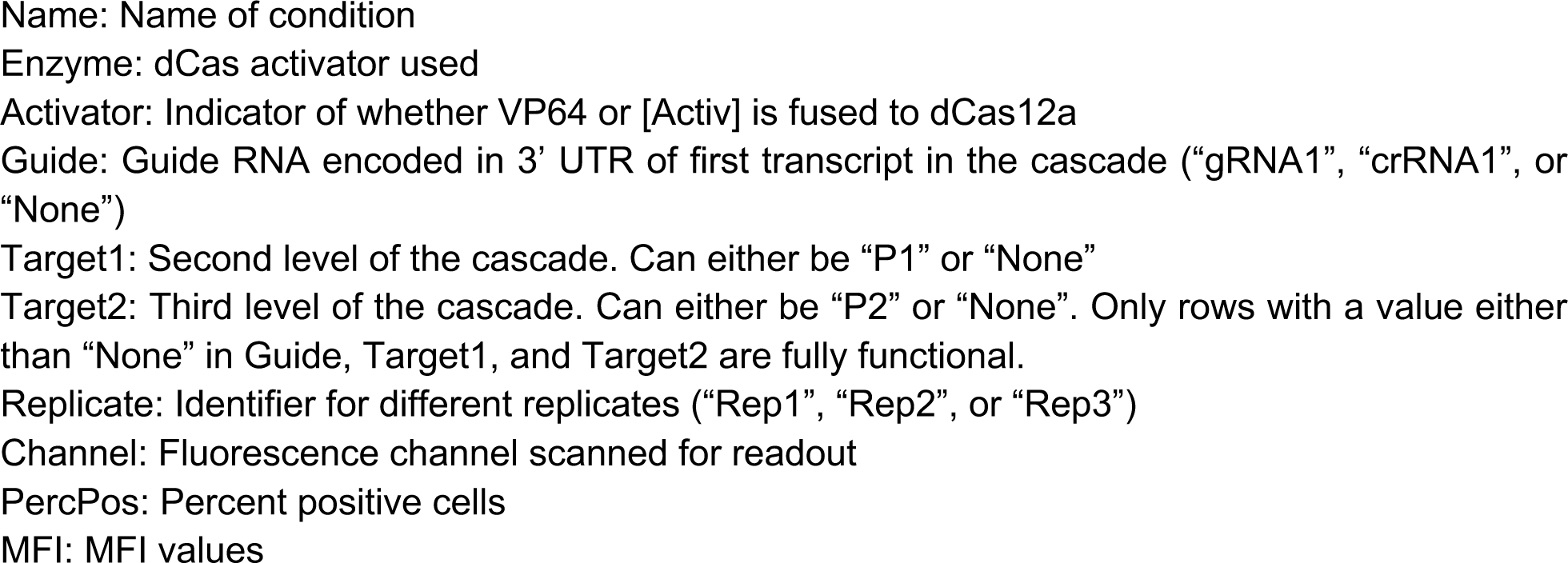
Three-level cascade (relevant to Figure 5)

## Notes

### Competing Interest Statement

The authors have declared no competing interest.

